# Coevolution of interacting proteins through non-contacting and non-specific mutations

**DOI:** 10.1101/2021.10.07.463098

**Authors:** David Ding, Anna G. Green, Boyuan Wang, Thuy-Lan Vo Lite, Eli N. Weinstein, Debora S. Marks, Michael T. Laub

## Abstract

Proteins often accumulate neutral mutations that do not affect current functions^1^ but can profoundly influence future mutational possibilities and functions^2–4^. Understanding such hidden potential has major implications for protein design and evolutionary forecasting^5–7^, but has been limited by a lack of systematic efforts to identify potentiating mutations^8,9^. Here, through the comprehensive analysis of a bacterial toxin-antitoxin system, we identified all possible single substitutions in the toxin that enable it to tolerate otherwise interface-disrupting mutations in its antitoxin. Strikingly, the majority of enabling mutations in the toxin do not contact, and promote tolerance non-specifically to, many different antitoxin mutations, despite covariation in homologs occurring primarily between specific pairs of contacting residues across the interface. In addition, the enabling mutations we identified expand future mutational paths that both maintain old toxin-antitoxin interactions and form new ones. These non-specific mutations are missed by widely used covariation and machine learning methods^10,11^. Identifying such enabling mutations will be critical for ensuring continued binding of therapeutically relevant proteins, such as antibodies, aimed at evolving targets^12–14^.

## Main Text

The ability of biological systems to maintain old functions and attain new ones after acquiring random mutations forms the substrate on which natural selection acts. It is unclear how this process is affected by neutral mutations that do not, on their own, change the function of the system, but can shape which future mutations are possible. Do these neutral mutations tend to restrict (Fig. 1a, left) or broadly expand the mutations that can subsequently arise (Fig. 1a, right)? In the absence of systematic studies, the identities and hidden potential of such neutral mutations in enabling subsequent mutational trajectories remain unclear.

**Figure 1:**
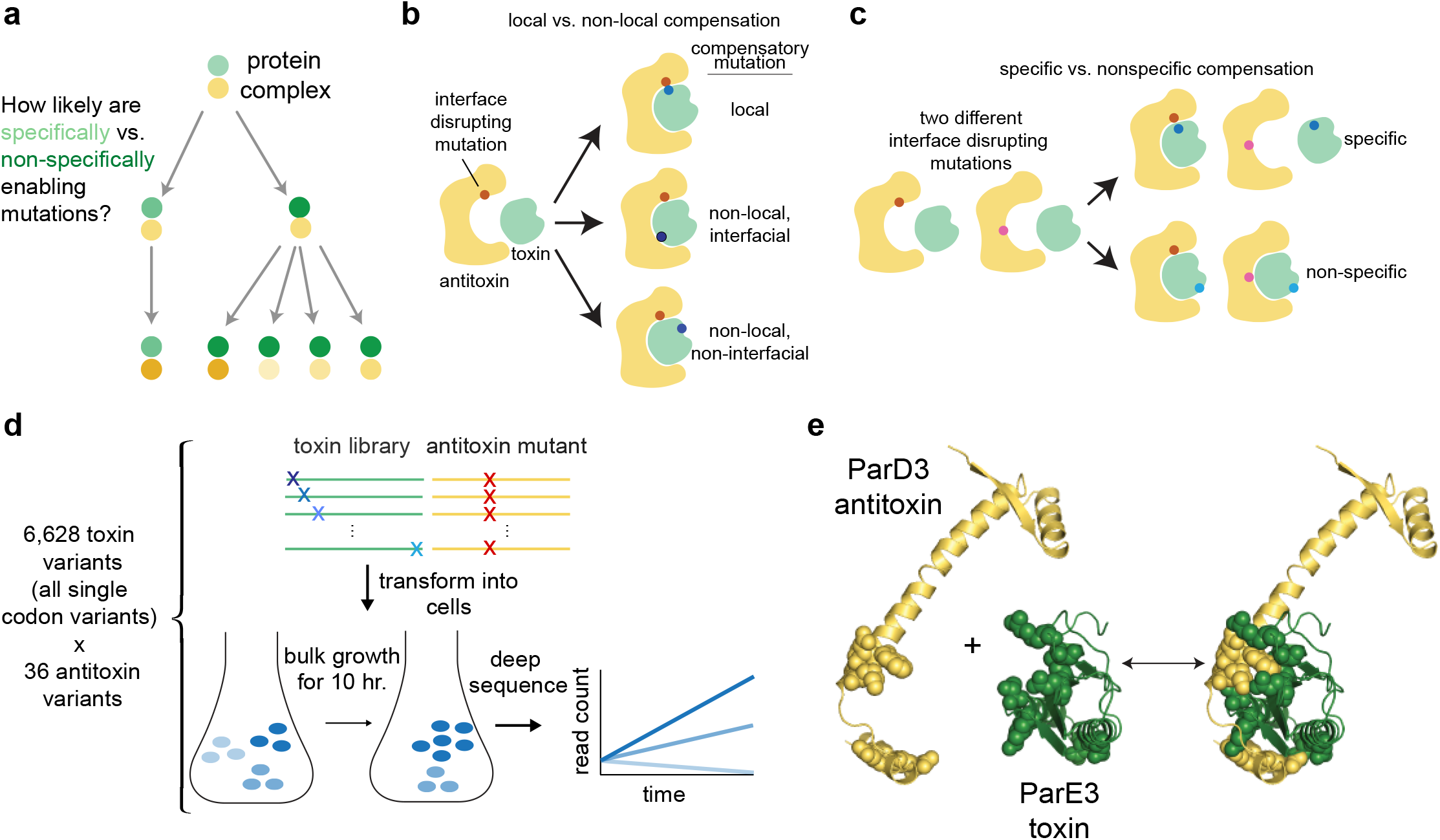
Comprehensive identification of neutral and enabling mutations for toxin-antitoxin ParD3/ParE3. **a**, Schematic of possible future mutational trajectories enabled by specific or non-specific neutral mutations. Yellow and green circles represent interacting proteins, shaded circles indicate mutated proteins, and arrows represent single mutations to sequences that retain binding. Specific mutations allow only particular subsequent mutations (left), whereas non-specific mutations enable tolerating many different subsequent mutations in the partner protein (right). **b-c**, Schematic examples of local vs. non-local (**b**) and specific vs. non-specific (**c**) compensatory mutations that rescue interface-disrupting mutations. Dots represent mutations. **d**, Schematic summary of experimental pipeline for identifying enabling mutations across the antitoxin-toxin interface. A library of all possible toxin single mutants is transformed into cells with a given, interface-disrupting mutation in the antitoxin (top). Cells are then grown in bulk and the abundance of each toxin variant over time is measured by sequencing. These changes are used to infer growth rates. Dots represent mutations. **e**, A single chain of the ParD3 antitoxin (yellow) in complex with the ParE3 toxin (green), from PDB:5CEG (right), with isolated antitoxin (left) and toxin (middle). The top 10 covarying positions are spacefilled.

For interacting proteins, several types of mutations can, in principle, enable tolerance to mutations that would have otherwise been disruptive (Fig. 1b-c). One is the mutation of a residue that contacts the disruptive mutation to directly restore the protein-protein interaction (Fig. 1b). The notion that interacting proteins evolve through such restricted and specific, complementary changes in contacting residues is suggested by analyses of amino-acid covariation in natural sequences, which can be used to predict protein structures and protein-protein interactions^15–20^ by identifying specific pairs of residues in close proximity^21–23^. Additionally, potent binding proteins have been engineered by mutating key interface residues^24–27^. However, some interface-disrupting mutations may be tolerated by mutations elsewhere in the partner protein, either along the interface or not, that indirectly restore the interaction (Fig. 1c). There are anecdotal examples, at least within proteins, of mutations that can only be tolerated in the presence of prior, enabling mutations at non-contacting positions^4–6,28–44^, and of mutations away from the interface regions of antibodies that affect antigen affinity and specificity^12–14^.

For virtually all protein-protein interactions, it is not yet known how many enabling mutations exist among the set of possible neutral mutations, nor is it known how close and how specific such mutations typically are to the disruptive mutations they enable^8,9^. Additionally, whether neutral, enabling mutations affect the evolvability of a protein - its ability to subsequently acquire new functions or binding partners - has not been systematically examined. A better understanding of how neutral mutations shape future mutational trajectories and protein binding partners promises to aid efforts to engineer protein interactions of clinical or therapeutic value, and enable better forecasting of fast evolving protein sequences.

To comprehensively study enabling mutations in protein coevolution, we examined a bacterial toxin-antitoxin system. The *Mesorhizobium opportunistum* antitoxin ParD3 normally binds and restrains the activity of its cognate, co-operonic toxin ParE3^21^. When co-expressed in *Escherichia coli*, ParD3-ParE3 form an inert complex and cells can grow. Any mutation that disrupts the interface will liberate toxin, which slows cell growth. Thus, cell proliferation provides a powerful, easy-to-measure readout of the ParD3-ParE3 interaction *in vivo* (Fig. 1d). Prior analyses of amino-acid covariation in ParD-ParE homologs identified residues in each protein that most strongly covary, map to the protein-protein interface, and can dictate interaction specificity of paralogs^21,22^ (Fig. 1e). To experimentally probe the possible mutational trajectories for ParD3-ParE3, we first used deep mutational scanning^45–48^ to identify all possible single point mutations in the toxin that are neutral and thus retain binding to the antitoxin and retain toxicity if produced alone. Then, we identified all possible mutations in the toxin that also enable it to tolerate interface-disrupting mutations in the antitoxin.

We find that the majority of such enabling mutations in the toxin can restore binding to many, and in some cases all, of the otherwise disruptive single mutations in the antitoxin. Hence, we refer to these as *non-specific suppressors.* These enabling mutations often arise far from the mutations they rescue, and some are even outside the protein-protein interface. If they arise before a disrupting mutation arises, these non-specific suppressors would stabilize the toxin-antitoxin interaction and thereby increase the robustness of this interaction to subsequent mutations. Notably, very few mutant pairs – a disruptive mutation and a suppressor mutation – correspond to the pairs identified in covariation analyses, suggesting that protein coevolution may often involve residues that are not directly contacting. We also demonstrate that non-specific suppressor mutations in the toxin enable it to bind substantially more antitoxins that have multiple mutations. Importantly, this includes an increased number of antitoxin variants that have the ability to also bind a different toxin. Thus, the enabling mutations in the toxin potentiate the evolvability of this protein-protein interaction by promoting binding to antitoxins with additional functions. Collectively, our findings highlight the potential of neutral, non-specific mutations in expanding the space of future mutational possibilities in protein coevolution and likely protein design.

## Results

### Deep mutational scanning reveals the mutational tolerance of the ParD3 antitoxin

To examine the mutational robustness of the ParD3-ParE3 (antitoxin-toxin) complex, we first built a library of 5,796 variants containing each possible codon at each position in the 93 amino-acid ParD3 antitoxin. This library encodes all 1,784 possible single amino-acid variants and 242 different synonymous versions of wild-type ParD3. The library was transformed into and expressed in cells harboring the wild-type toxin ParE3. If a cell has an antitoxin variant that can bind and neutralize the toxin, it will grow and proliferate in the population over time; conversely, if a cell has an antitoxin variant that disrupts the interface and liberates toxin, it will not grow and eventually be lost from the population. The growth rates of individual variants were assessed in pooled cells, using deep-sequencing to measure the change in variant frequency over 10 hours, which is ~6 generations for cells harboring wild-type ParD3 and ParE3. To infer the effect of each substitution from sequencing read counts, including their uncertainties, we developed a hierarchical Bayesian error model that considers sampling noise of reads and information from synonymous variants and replicate experiments (see Methods). The inferred growth rates were highly correlated with independently measured growth rates (Pearson r=0.94; Extended Data Fig. 1a), and the change in variant frequency over time was highly reproducible between biological replicates (Pearson r=0.92, Extended Data Fig. 1b).

For each variant in the antitoxin library we calculated Δgrowth-rate - the difference in number of doublings of the mutant, AT*, compared to wild-type antitoxin, AT (Fig. 2a). For synonymous variants of the wild-type antitoxin, the Δgrowth-rate values were tightly and symmetrically distributed around 0, as expected. For variants with a stop codon that would produce a non-functional antitoxin, the Δgrowth-rate values had a mean of −5.2, also as expected. For all other variants, the Δgrowth-rate values had a mean of −0.5, with > 98% of all variants > −2.5. Only 32 amino-acid variants produced Δgrowth-rate values < −2.5 indicating that few single substitutions severely disrupt the toxin-antitoxin interface.

**Figure 2:**
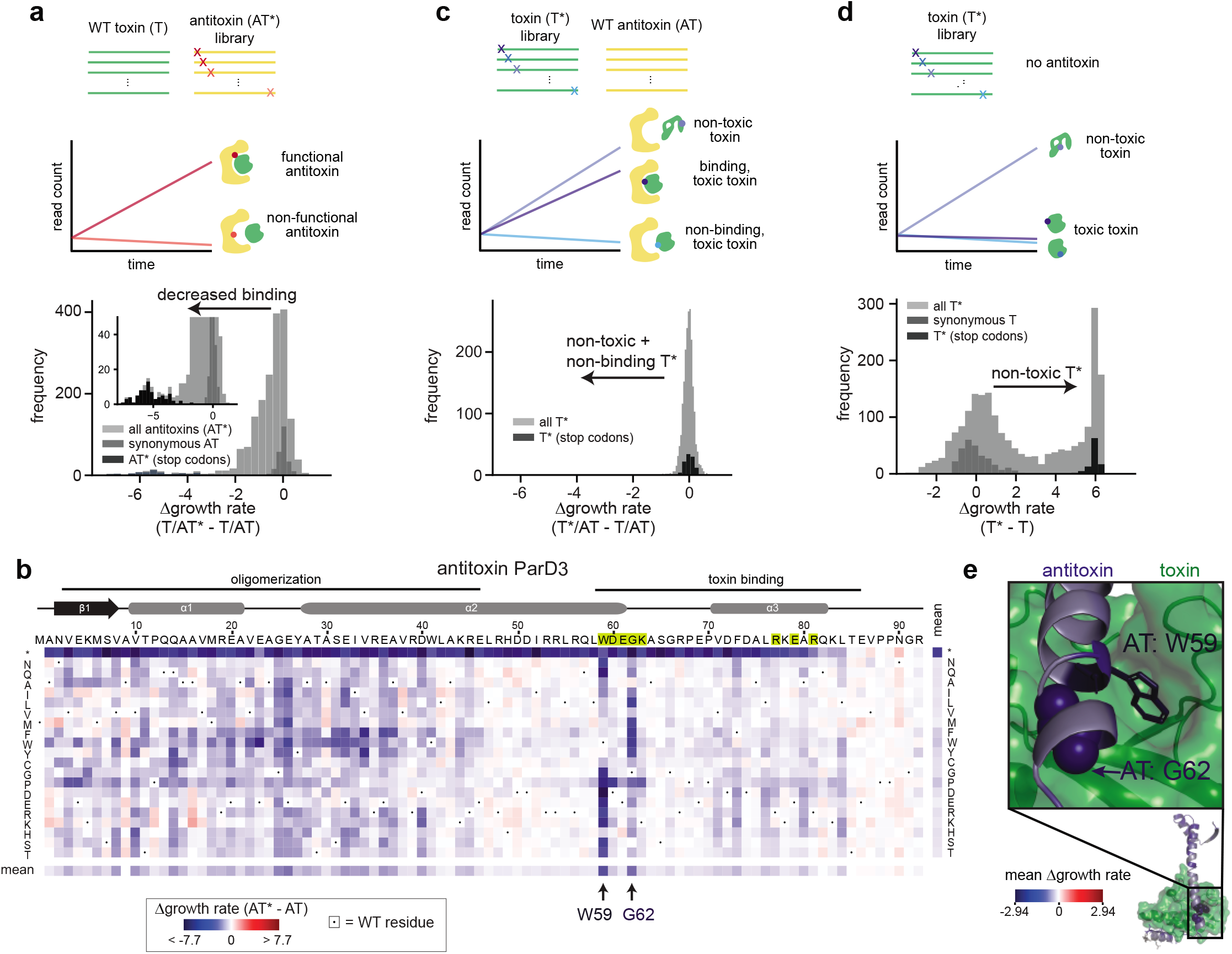
Deep muational scanning reveals mutational tolerance and interface-disrupting substitutions in ParE3-D3. **a**, Comprehensive single mutant scan of all possible antitoxin variants. The histograms indicate the change in growth rate for each mutant (with a blow-up histogram inset), relative to the wild-type antitoxin or toxin, with the greyscale-coded categories indicated. **b**, Heatmap of Δgrowth-rate values for each possible antitoxin single mutant showing mutations that disrupt toxin neutralization (blue). The antitoxin-antitoxin oligomerization and antitoxin-toxin binding regions are indicated above. Top 8 positions that covary with positions in the toxin are shaded in yellow on the primary sequence. The substituted residue (or stop codon indicated with *) is listed on the far left and far right. Mean effects for each row and column are shown below and to the right. **c,d**, Same as (a) but for all toxin variants in presence of wild-type antitoxin (**c**) or absence of antitoxin (**d**). **e**, Structure of ParD3-ParE3 (PDB ID: 5CEG) highlighting antitoxin residue W59 with its pocket in the toxin and antitoxin residue G62 where single substitutions disrupt the interaction most. The mean effect at each position (see panel b) in ParD3 is color-coded, as indicated, on the structure.

To examine the position-wise effect of amino-acid substitutions, we generated a 21 x 92 heatmap showing the Δgrowth-rate value for each possible substitution at each position in ParD3 (Fig. 2b). This map revealed an apparent periodicity within a-helices with residues on the face of an a-helix that contacts the toxin exhibiting negative mean Δgrowth-rate values (Extended Data Fig. 2a), whereas solvent-exposed residues on the opposite face exhibited more mutational tolerance. Helix-breaking proline substitutions did not follow this periodicity and were disruptive at most positions in a-helices. Aside from these prolines, the substitutions disruptive to growth rate were generally clustered in two regions. One was the N-terminal half of the antitoxin, which mediates oligomerization and likely promotes overall stability of the ParD3-ParE3 complex (Extended Data Fig. 2b). The second region involves the C-terminal end of α2 and a3, the elements of ParD3 that bind the toxin (Fig. 2b).

The most disruptive substitutions in ParD3 arose at just two positions, tryptophan-59 and glycine-62 (Fig. 2b). These positions strongly covary with nearby residues (< 6 Å minimum atom distance) in the toxin as measured by EVcouplings^18^ (Extended Data Fig. 3) and are critical for antitoxin binding specificity^21^. For position 59, substituting tryptophan with another aromatic residue or some hydrophobic residues had relatively modest effects, with all others leading to substantial defects (Δgrowth-rate < −2.6). For position 62, substituting glycine with anything other than alanine, valine, or serine severely disrupted growth. Notably, W59 and G62 are at the very C-terminus of α2 in the antitoxin and both are in close proximity to the toxin. The side chain of W59 inserts into a snug hydrophobic pocket on the toxin and G62 is tightly packed against the toxin (Fig. 2e). The substitutions at positions W59 and G62 that severely diminish growth rate may either disrupt the toxin-antitoxin interface or trivially unfold the antitoxin. Below we provide evidence for the former.

### Deep mutational scanning reveals the mutational robustness of the ParE3 toxin

Next, we identified all possible single substitutions in the toxin that maintain both toxin-antitoxin binding and toxicity of the toxin. We performed a deep mutational scan of the 103 amino-acid toxin ParE3 (Fig. 2c), using a library of all possible 6,426 single-codon variants (2,040 amino-acid variants). The distribution of Δgrowth-rate values in the presence of wild-type antitoxin (*i.e*. the growth rate of each toxin variant, T*, relative to the wild-type toxin, T) was narrowly centered around 0. Thus, every possible single substitution either retains binding to the antitoxin or causes the toxin to lose toxicity, possibly through unfolding.

To identify mutations that allowed cells to grow by disrupting the toxicity of the toxin rather than retaining antitoxin binding, we repeated the experiment but in cells lacking the antitoxin (Fig. 2d, Extended Data Fig. 4a-c). In this case, unfolded or nonfunctional toxin variants permit growth, whereas properly folded and functional toxins do not. For variants containing a synonymous mutation (n=278), the Δgrowth-rate values were symmetrically centered around 0, while variants harboring a stop codon, which are presumably non-functional, exhibited high Δgrowth-rate values (> 5) indicating significantly faster growth than cells expressing wild-type toxin. For all other variants, the distribution was bimodal, with one mode at 0 and the other at ~6. The set nearest to 0 are those that minimally disrupt toxin folding and activity. We performed this experiment using 7 different induction conditions for the toxin, and then identified the 310 ‘most toxic’ variants at 65 residue positions whose ability to inhibit growth rate was consistently comparable to the wild-type toxin (see Methods, Extended Data Fig. 4c). We also used a less stringent cutoff to define a set of 781 ‘toxic’ mutants at 91 residue positions that are comparable to wild-type toxin under 4 fully inhibiting induction conditions (Extended Data Fig. 4c).

Substitutions that ablated ParE3 toxicity (and hence have high Δgrowth-rate values) were particularly pronounced in β2, β3, and α3, which have low solvent accessibility, suggesting that they simply unfold the toxin (Extended Data Fig. 4d-f). Notably, for toxin positions that strongly covary with positions in the antitoxin and that make direct contact in the co-crystal structure, there were few substitutions that produced high growth rates indicative of unfolding (Extended Data Fig. 4g). Substitutions at these positions also retained their neutralization by the antitoxin (Extended Data Fig. 4h). Taken together, our results indicate that the interface residues of the ParE3 toxin are highly robust to mutations. This robustness does not result from overexpression of the antitoxin, as our experiments were performed with minimally neutralizing levels of antitoxin (Extended Data Fig. 5).

### Suppressor scanning reveals enabling mutations distant from interface-disrupting mutations

We next sought to systematically assess which mutations in the toxin enable binding to otherwise interaction-disrupting mutations in the antitoxin. To this end, we performed a ‘deep suppressor scan’, by screening all toxin single mutations for their ability to rescue a disruptive mutation in the antitoxin. We chose 36 single antitoxin substitutions, spanning the measured range of effects, mostly at positions that (i) strongly covary with positions in the toxin and (ii) that involve amino acids commonly found at those positions in ParD antitoxin homologs (see Methods). We examined each antitoxin mutant + toxin library (36*2,040 = 73,440 amino-acid variant pairs) at two different induction levels of antitoxin: the minimal concentration needed to neutralize wild-type toxin, as above, and a slightly lower concentration of antitoxin such that the toxin is almost but not fully neutralized, which enables more sensitive detection of rescuing mutations in the toxin. Of the 36 antitoxin mutants examined, 9 were deleterious at the higher antitoxin concentration, and 12 at the lower concentration. We focused on the 9 most deleterious antitoxin mutants, but found that including data from all 36 in our model (see below) improved our assessment of the effect of each toxin mutation.

As an example, we first considered the antitoxin mutant ParD(W59T), which strongly disrupts the ParD3-ParE3 interface (Fig. 3a, Fig. 2b). Toxin variants with positive Δgrowth-rate values in this antitoxin background either increase binding to the W59T antitoxin mutant, or simply disrupt the toxicity of the toxin. Considering only the 310 ‘most toxic’, neutral mutations defined above that maintain the toxicity of the toxin, we identified 11 mutations in the toxin that significantly and substantially improved the growth rate of cells harboring the ParD3(W59T) mutant (p < 0.0001 and Δgrowth-rate values > 0.5, see Methods; Fig. 3b, Extended Data Fig. 6). We mapped these 11 rescuing mutations onto the ParD3-ParE3 co-crystal structure (Fig. 3c), finding that they were distributed throughout the toxin and many were > 10 Å away from the W59 residue in the antitoxin (Fig. 3d).

**Figure 3.**
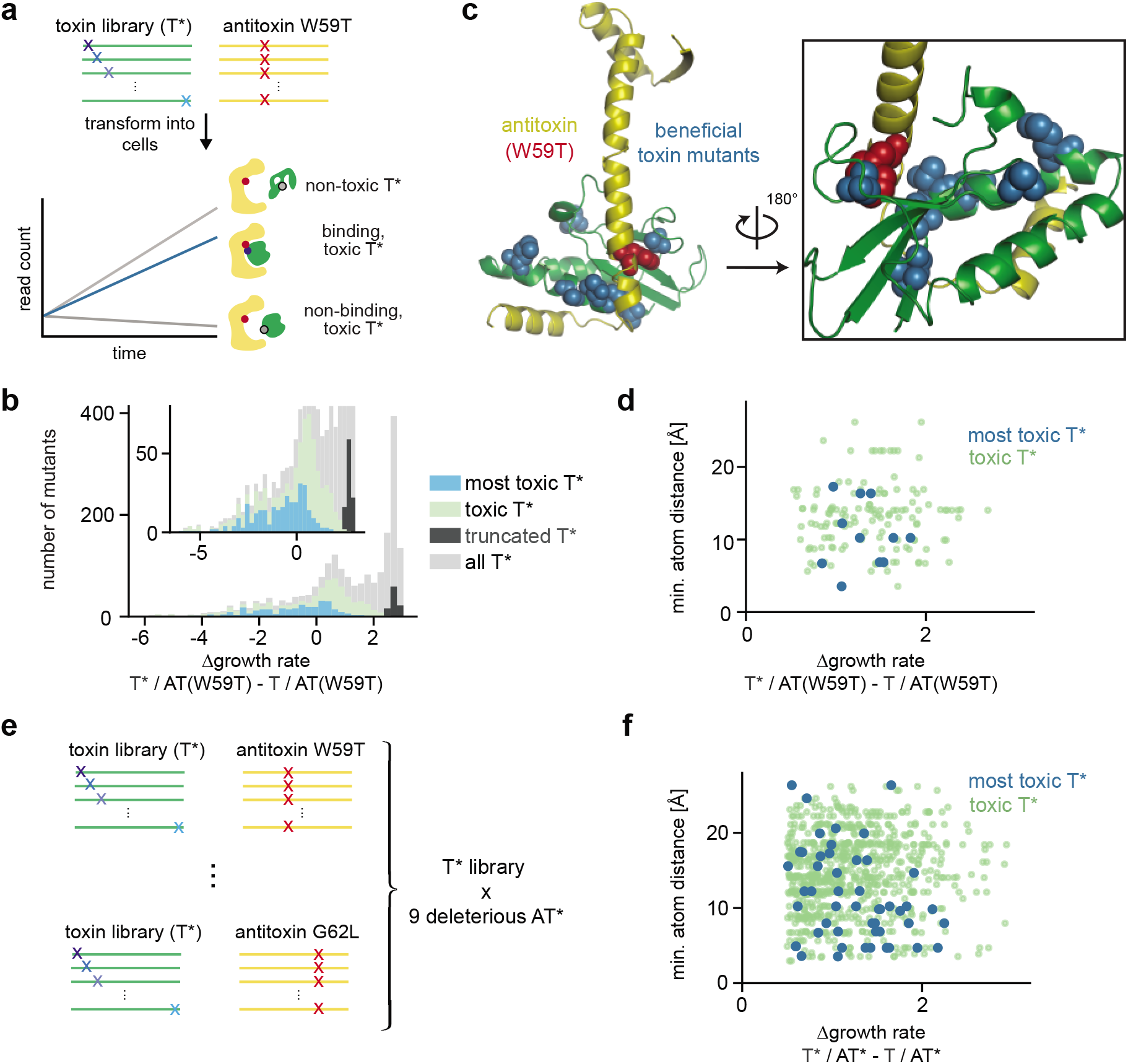
Beneficial, interaction-restoring mutations can be far from the deleterious mutation they rescue. **a**, Schematic overview of ‘suppressor scanning’. Cells expressing antitoxin ParD3(W59T) and a library of all possible toxin single substitutions were grown and analyzed as in Fig. 1f to identify toxin variants (T*) that can rescue the growth defect of ParD3(W59T). **b**, Distribution of growth rates for each toxin variant (T*) relative to the wild-type toxin (T) when co-expressed with antitoxin ParD3(W59T) reveals toxin variants alleviating the growth defect of the antitoxin W59T mutation (with a blow-up inset). Various categories of toxin variants are color-coded as indicated (right), including toxin variants that maintain toxicity at different thresholds (blue: 310 most toxic toxin variants, green: 781 toxic toxin variants). **c**, The significantly beneficial toxin variants for the deleterious antitoxin W59T (blue spacefilled, from set of most toxic toxin variants) are distributed across the toxin in the ParD3-ParE3 structure (PDB ID: 5CEG). Red indicates the deleterious antitoxin residue W59. **d**, Plot of distance between W59 in ParD3 and each significantly beneficial toxin mutant from the set of toxic (green) or most toxic (blue) toxin variants vs. effect size of rescue. **e**, Schematic indicating that all toxin single mutants were screened against 9 deleterious antitoxin mutants (G62L/D/Y, W59A/L/T/V, K63D,F73K). **f**, Same as (d) but for significantly beneficial toxins from all 9 suppressor scans.

We repeated the same analysis for the other 8 deleterious antitoxin mutations at the higher antitoxin concentration (Fig. 3e, Extended Data Fig. 7). In the pooled data, we detected 51 pairs of mutations in which the toxin variant significantly and substantially alleviates the growth defect of a deleterious antitoxin mutation (Fig. 3f). For these pooled data, there was no strong correlation between the magnitude of rescue by toxin mutations and their distances to the position mutated in the antitoxin (Fig. 3f). Similar results were seen for the 12 deleterious mutations at the lower antitoxin concentration (Extended Data Fig. 8). We conclude that there are many possible mutations that can relieve the deleterious effect of a given antitoxin mutation and that such mutations can arise throughout the toxin, not simply through local, directly compensating mutations.

### Many enabling mutations in the toxin non-specifically tolerate many different antitoxin mutants

Next, we sought to assess whether enabling toxin mutations are non-specific or specific, *i.e*. whether a toxin mutation increases binding irrespective of the deleterious antitoxin mutation present, or allows binding only to particular deleterious antitoxin mutations. To do this quantitatively, we built a model, similar to previous models^49–58^, that tries to explain all observed single and double mutant growth rates as a sum of non-specific, independent (but unobserved) single substitution effects passed through a global nonlinearity (see Methods, Extended Data Fig. 9). This model explained 89% of the observed growth rate deviations (Extended Data Fig. 9b), with inferred mutation effects highly robust to expression conditions (Extended Data Fig. 9h), and allowed us to calculate the expected growth rate of each double mutant if the toxin and antitoxin substitutions act independently and non-specifically. We defined a toxin-antitoxin double mutant pair as specific if it showed a significant and substantial positive deviation from its double mutant expectation (>2-fold change in growth rate, and p < 0.0001; Fig. 4a,c; Extended Data Fig. 10), and as non-specific if the observed growth rate was close to the expected rate (Fig. 4b,c; Extended Data Fig. 10). We then called a particular toxin mutation as specific if it showed a positive deviation in any antitoxin background, and conversely called a toxin mutation non-specific if it produced growth rates close to the expectation in all antitoxin backgrounds.

**Figure 4.**
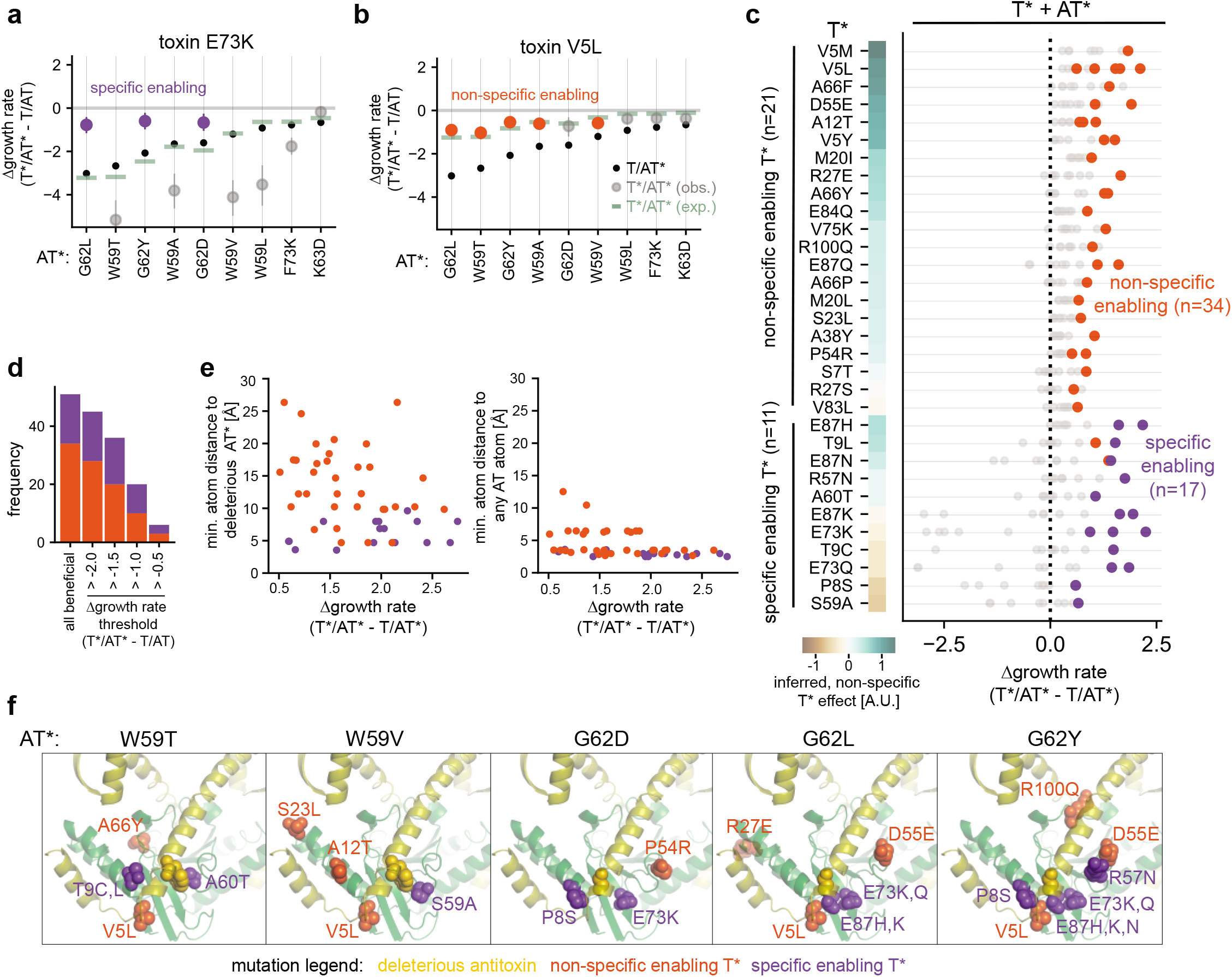
Non-specific enabling mutations outnumber specific mutations, and can be far from the deleterious mutation as well as the interface. **a-b**, For a specifically enabling toxin mutation E73K (**a**) or non-specifically enabling toxin mutation V5L (**b**), the growth-rate relative to the wild-type pair T/AT (Δgrowth rate) is shown when combined with each antitoxin variant indicated on the x-axis (large dots represent T*/AT*; error bars indicate 95% posterior highest density interval). The Δgrowth rate for each AT* combined with wild-type T is shown (small black dots) along with the Δgrowth rate for T*/AT* expected under the independent, nonlinear model (green lines). Purple and orange indicate T*/AT* pairs (T*/AT*) where the toxin substitution is specific or non-specific, respectively. **c**, For each toxin variant (n=32) beneficial to at least one deleterious antitoxin mutant (51 pairs), the plot shows their effect size of rescue (Δgrowth rate) combined with each of the 9 deleterious AT*. The effect of each T* inferred by the nonlinear model is indicated on the heatmap. **d**, The number of non-specific and specific pairs of rescue at different Δgrowth rate thresholds relative to the wild-type T/AT. **e**, Minimum atom distance of rescuing toxin mutation to deleterious antitoxin mutation (left) or to any antitoxin residue (right) vs. effect size of rescue (Δgrowth rate). For color codes of toxin-antitoxin double mutant pairs, see panels (a)-(b). **f**, Specific (purple) and non-specific (orange) enabling toxin mutations in each antitoxin mutation background they rescue are shown (spacefilling) on the ParD3-ParE3 structure (PDB ID: 5CEG). Antitoxin is yellow; toxin is green.

The 51 mutation pairs in which the growth defect of a deleterious antitoxin mutation is alleviated by a toxin mutation, include 32 different toxin variants, and we found that 21 of these toxin variants act non-specifically, outnumbering the 11 specific toxin variants (Fig. 4c, Extended Data Fig. 11a-b). These 51 mutant pairs involved any toxin mutation that significantly improved the growth rate of an antitoxin mutant, regardless of how close the double mutant growth rate was to the wild-type toxin-antitoxin pair. We also examined only those pairs in which the double mutant had a growth rate close to the wild-type toxin-antitoxin pair. At each threshold considered, the number of non-specific, beneficial toxin mutations was the same or greater than the number of specific beneficial mutations (Fig. 4d). These conclusions were even more pronounced under the ‘low antitoxin’ expression conditions (Extended Data Fig. 11c-e).

To probe the spatial distribution of specific and non-specific toxin mutants, we used the ParD3-ParE3 co-crystal structure to assess whether each lies at the protein interface and to measure the distance of each from the antitoxin mutation they rescue (Fig. 4e-f). Specific rescuing toxin mutations were all within 10 Å of the antitoxin mutation they rescued with nearly half within ~5 Å. However, these pairs did not necessarily arise from biochemically obvious compensation, such as charge swapping or preservation of size complementarity. In contrast to the specific rescuing mutations, the non-specific beneficial toxin substitutions were mostly far (> 6 Å when considering minimum atom distance) from the antitoxin substitutions they rescued (includes 31 of the 34 pairs involving a non-specific rescuing mutation, which involves 20 of the 21 non-specific toxin mutations) (Fig. 4e-f, Extended Data Fig. 11g), with 8 of 21 non-specific toxin mutations at positions more than 6 Å from any antitoxin residue indicating that they are not toxin-antitoxin interface residues (Extended Data Fig. 11f, Table S1, File S1). Taken together, our analyses indicate that non-specifically enabling mutations are more frequent than specifically enabling mutations and that many of these non-specific toxin mutations arise at sites spatially distant from the mutation they rescue.

### Analyses of natural sequence variants cannot predict enabling mutations

We next asked whether the outcome of our ‘deep suppressor scan’ could have been predicted from the features of naturally occurring homologs of ParD3-ParE3, which show strong covariation scores, even compared to other complexes^15^ (Fig. 1e; Fig. 5a,b). The top covarying positions involve nearby pairs of interface residues (28 of top 29 covarying pairs of residues are within 6 Å minimum atom distance, Extended Data Fig. 3). However, the distribution of covariation scores for position pairs identified as enabling in our suppressor scan was indistinguishable from that of randomly selected pairs (Fig. 5c). There was also no correlation between the effect of each toxin mutation inferred in our nonlinear model and the frequency at which the mutant amino acid is found at that position in natural sequences (Fig. 5d). Similarly, pairwise frequencies or enrichments thereof were not predictive of beneficial pairs of residues, with more than half (29/51) of the beneficial toxin and deleterious antitoxin variant pairs not observed in natural sequences (Extended Data Fig. 12).

**Figure 5.**
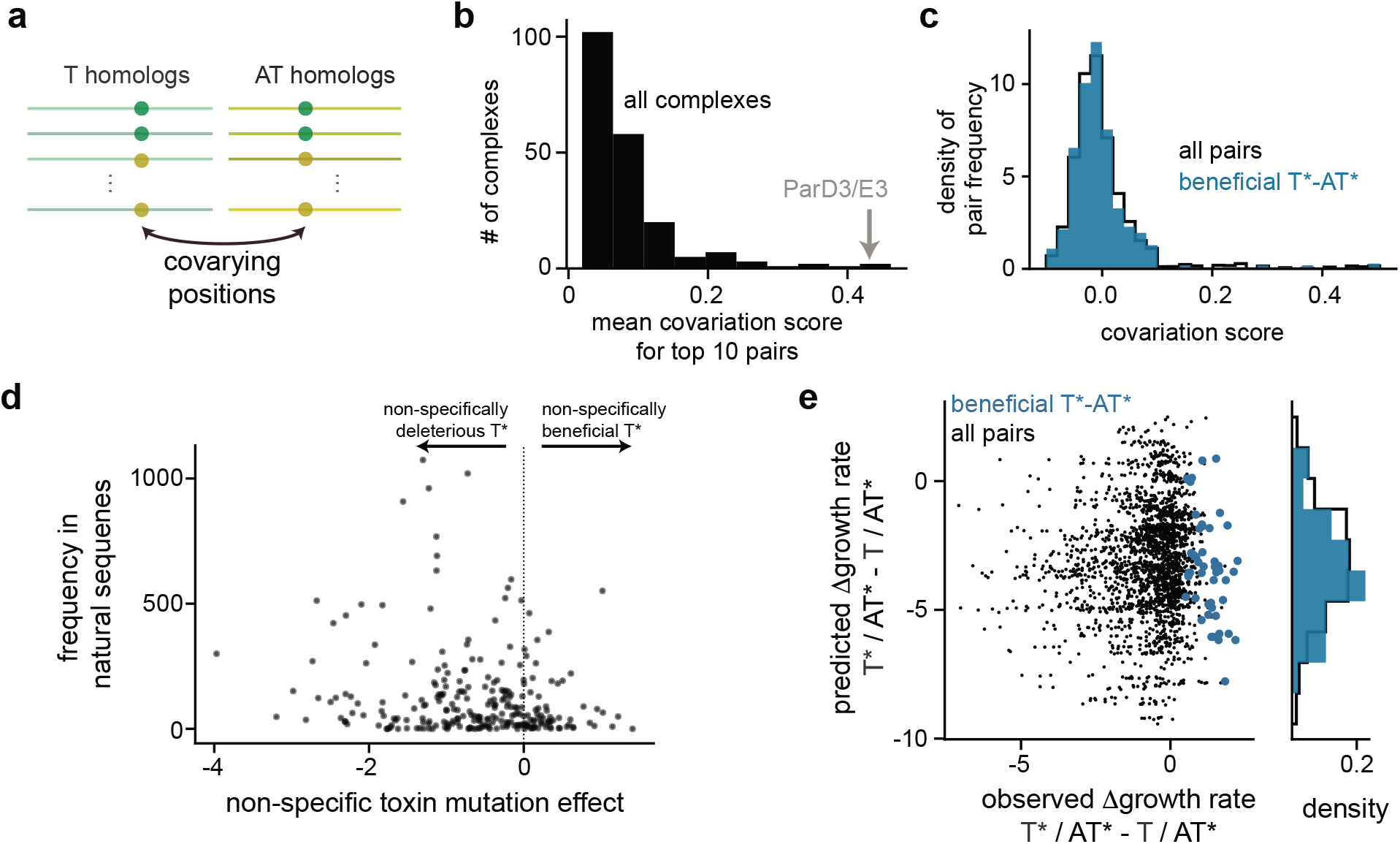
Natural sequences and models trained on these provide insufficient information to predict enabling mutations. **a**, Schematic of identifying covarying pairs of residues in natural sequences. **b**, Mean covariation score of top 10 pairs of residues in ParD/ParE homologs (grey) compared to ~350 complexes with covariation signal (black). **c**, Distribution of covariation scores for beneficial pairs of residues (blue, see Fig. 2e) is similar to the null distribution for all possible pairs of positions (black). **d**, Amino-acid mutant frequency in toxin homologs at a particular site is not correlated with the non-specific enabling effect of that amino acid variant, as inferred from suppressor scanning. **e**, Predicted vs. observed rescue (Δgrowth rate of each double mutant (T*/AT*) relative to the antitoxin single mutant effect (T/AT*)). Predictions are made using EVmutation. Blue indicates observed significant rescue.

We also found that models trained on the sequence alignment (EVmutation^10^, variational autoencoder^11^) could not predict which toxin mutations would alleviate the growth defect of a particular antitoxin mutation, with no correlation between the measured Δgrowth-rate values and the scores produced by these models (Fig. 5e, Extended Data Fig. 13). Notably, the homologs in our alignment differ, relative to our ParD3-ParE3 complex, at 80% of positions, on average (Extended Data Fig. 12i). These natural sequences may effectively be too sparse of a dataset to predict the immediately available compensating mutations, either specific or non-specific, available in a particular sequence background.

### Non-specific suppressors expand subsequent mutational trajectories for evolving additional binding functions

The non-specific enabling mutations identified in the ParE3 toxin promote tolerance to many different single mutations in the antitoxin. To ask whether they also expand subsequent mutational trajectories containing multiple antitoxin mutations, we performed a ‘combinatorial mutation scan’ using a previously developed library containing all 8,000 possible combinations of residue variants at 3 interface positions in the ParD3 antitoxin critical to binding specificity^22^ (D60, K63, E79) (Fig. 6a).

**Figure 6.**
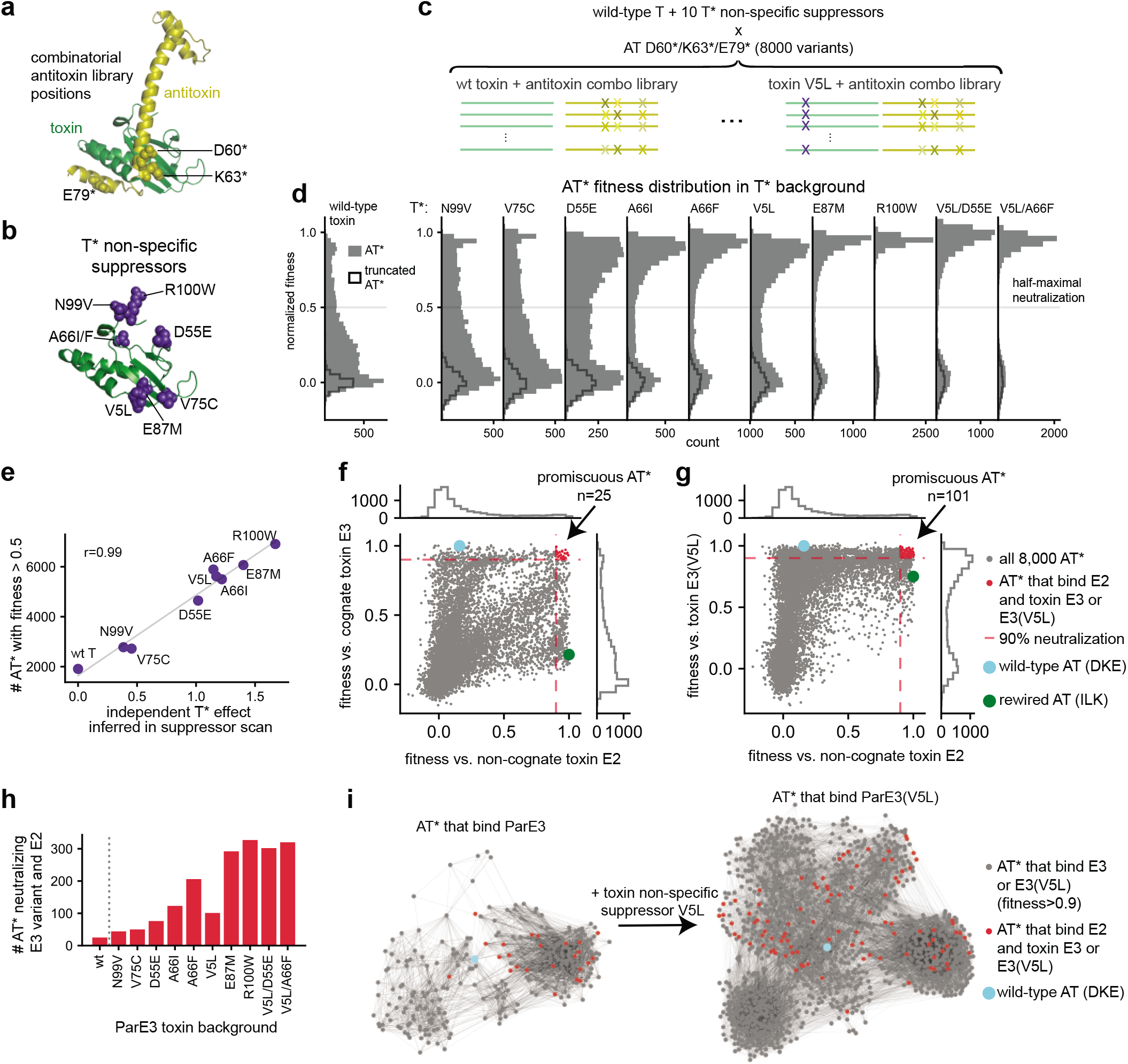
Non-specifically enabling mutations expand mutational paths to maintain old and evolve new interactions. **a-c**, For the antitoxin ParD3, a library of all 8,000 possible variant combinations at three key specificity-determining residues (shown on the structure of ParD3-ParE3 (PDB ID: 5CEG) in (**a**)) was screened for binding to wild-type toxin ParE3 or ParE3 harboring non-specifically enabling variants (purple) shown in (**b**). Schematic of the experiment is shown in (**c**). **d**, The fitness distribution of 8,000 antitoxin variants screened against the wild-type toxin ParE3 (left) or 10 different variants of ParE3 (right) reveals that non-specifically enabling mutations in the toxin allow binding to more combinatorial antitoxin variants than the wild-type toxin. **e**, Correlation of inferred independent effect from suppressor scan (see Fig. 2) with the number of combinatorial antitoxin variants neutralized above half-maximal for each toxin single variant. **f,g**, Scatterplots showing fitness values for 8,000 antitoxin variants (dots) screened against the wild-type toxin ParE3 (y-axis) and the non-cognate toxin ParE2 (x-axis) (**f**), or against toxin ParE3 with a non-specifically enabling mutation (ParE3(V5L)) (y-axis) and ParE2 (x-axis) (**g**). Various classes of AT* are color-coded as shown on the right, including the AT* variants that have gained the ability to bind the non-cognate ParE2 (red). **h**, Differences in the number of promiscuous antitoxin variants that neutralize both the non-cognate toxin ParE2 as well as the wild-type type toxin ParE3 (left) or a toxin variant harboring a non-specific suppressor indicated on the x-axis (fitness > 0.9). **i**, Force-directed graph of antitoxin variants that bind toxin ParE3 with (left) or without (right) the non-specifically enabling mutation V5L. Nodes represent individual antitoxin sequences (fitness >0.9), edges correspond to single mutational steps. Color codes as in panel f+g.

We picked 8 of our non-specific suppressors in the toxin and two double mutants that combine two of these suppressors (Fig. 6b). We verified that each of these 10 toxin variants are as, or almost as, toxic as wild-type toxin (Extended Data Fig. 14). We then assayed each, and the wild-type toxin, for neutralization by the combinatorial library of 8,000 ParD3 antitoxin variants (Fig. 4c, Extended Data Fig. 15). In each case, we calculated a normalized fitness between 0 (comparable to a truncated antitoxin) and 1 (comparable to wild-type antitoxin).

Notably, each non-specific enabling mutation in the toxin led to a substantial increase in the number of antitoxin variants that could neutralize it (Fig. 6d). For the wild-type toxin ParE3, less than 20% of antitoxin variants achieve half-maximal neutralization (fitness > 0.5). In contrast, each non-specific rescuing mutation led to many more neutralizing antitoxin variants, often with > 60% of the library exhibiting fitness values > 0.5. The independent effect of each toxin mutant inferred from our initial suppressor scan was almost perfectly correlated with the number of antitoxin variants that could bind (Fig. 6e). We conclude that the global, non-specific mutations identified in our suppressor scan expand the subsequent mutational robustness of the toxin-antitoxin complex.

Whereas the wild-type antitoxin ParD3 can only neutralize its cognate toxin ParE3, we identified 25 promiscuous variants in the antitoxin combinatorial library that could also interact with the non-cognate toxin, ParE2^21,22^, which shows ~40% sequence identity with toxin ParE3 (fitness > 0.9, Fig. 6f; for other cutoffs, see Extended Data Fig. 16). We asked whether the non-specific enabling mutations in ParE3 increased the number of promiscuous antitoxin variants. Indeed, the number of antitoxin variants that could neutralize both ParE3(V5L) as well as ParE2 was 101, a 4-fold increase relative to the number that neutralize wild-type ParE3 and ParE2 (Fig. 6f-g). All of these promiscuous antitoxin variants are accessible from the wild-type ParD3 antitoxin via trajectories comprising single mutational steps (Fig. 6i, Extended Data Fig. 17). Even larger increases in the number of promiscuous antitoxin variants were obtained when considering other toxin ParE3 global suppressor variants (Fig. 6h, Extended Data Fig. 16). We conclude that the non-specific suppressors identified in ParE3 both improve its ability to maintain an interaction with the wild-type partner ParD3 and promote the evolution of new toxin-antitoxin interactions.

## Discussion

Collectively, our results indicate that interacting proteins may coevolve by first acquiring neutral, non-specific mutations in one protein that dramatically expand the number of mutations tolerated in the partner that would have otherwise disrupted the interaction. These enabling mutations are often found at positions far from the disruptive substitutions they promote tolerance to, and many lie outside of the protein-protein interface. These mutations not only promote maintenance of the specific cognate interaction, but expand the number of subsequent mutational trajectories that include partners with additional functions. Collectively, these findings contrast with models of coevolution driven mainly by directly contacting, specifically compensating pairs of mutations as it involves distant residues and because it does not invoke a broken, or disrupted, intermediate state. The non-specific mutations we identified, in fact, allow the inverse: a neutral, non-specifically enabling mutation could occur first, and then permit the mutation in its partner that would otherwise have disrupted the interface. Although anecdotal examples of non-specific suppressors have previously been reported within proteins^4,5,28–43,59,60^ and some between proteins^12–14^, it had been unclear in the absence of systematic studies how likely non-specific vs. specific suppressors are to arise and whether each class of suppressors would map to directly contacting pairs of residues or not^9^.

Our systematic ‘suppressor scan’ identified, for ParD3-ParE3, all possible compensatory mutations after introducing a handful of deleterious mutations. The 21 non-specific suppressors we identified arose at 15 different positions in the toxin, and are mostly far from the deleterious antitoxin substitution they rescue, sometimes without contacting the antitoxin at all. The mechanistic basis of these 21 non-specific suppressors identified in ParE3 is not yet clear. Some of these non-specific mutations may promote binding by forming new points of favorable interaction (Table S1, File S1), thereby enhancing complex formation and tolerance to subsequent mutations. They may also promote or stabilize existing points of interaction. In either case, the ‘excess’ binding energy may then permit a wide range of subsequent mutations that would have otherwise destabilized the complex. Such a model has also been proposed for non-specific suppressors of destabilizing mutations within proteins^4,43^. The non-specific toxin mutations likely increase binding to both wild-type and mutant antitoxins. Alternatively, the non-specific mutations could increase binding only to mutant antitoxins while retaining similar binding to the wild-type antitoxin, but such a scenario is less parsimonious as it invokes multiple specific mutation dependencies and is less biochemically plausible, especially for the non-specific mutations that do not lie on the ParD3-ParE3 interface.

Our strategy for identifying non-specific suppressors could be valuable in the design of therapeutic proteins aimed at an evolving target. For instance, systematically finding and engineering distal, non-specific suppressors of different magnitude into antibodies could help render them broadly neutralizing and tolerant to subsequent mutations in the target antigen. Examples of such mutations have been reported for certain broadly neutralizing antibodies^12,14^. Notably, the non-specific suppressors we identified were not predictable based on naturally occurring homologs. Although ParE-ParD homologs exhibit strong covariation and natural sequence homolog-based models are powerfully able to predict protein and protein complex structures, our results indicate that such models are currently insufficient to accurately predict the immediate mutational trajectories of our proteins. One explanation is that the covariation signal measured in sequence alignments may represent an average signal across all homologs found in nature, but that the exact mutational steps possible for each extant sequence are highly idiosyncratic. If so, systematic experimental methods, such as the suppressor scan performed here, will be critical to future protein engineering efforts.

## Acknowledgements

We thank members of the Laub and Marks labs, Alexandra Batchelor, Conor McClune, John Ingraham, Armin Schoech, and Ivana Cvijovic for helpful discussions. We thank Andrew Murray, Nicholas Gauthier, Tatsuo Okubo, Sam Sinai, and Nour Youssef for feedback on the manuscript, and Michael Stiffler for sharing protocols before publication. This work was supported by the Howard Hughes Medical Institute (M.T.L.), National Institutes of Health grant R01CA260415 (D.S.M.), Chan Zuckerberg Initiative CZI2018-191853 (D.S.M.), Ashford PhD fellowship (D.D.), Boehringer Ingelheim Funds PhD fellowship (D.D.), Fanny and John Hertz Fellowship (E.N.W.), National Institutes of Health NLM Training Grant T15LM007092 (A.G.G.), National Institutes of Health grant T32GM007753 (T.V.L.), Jane Coffin Childs Memorial Fund for Medical Research fellowship (B.W.) and National Institutes of Health grant K99GM135536 (B.W.).

## Author contributions

D.D., D.S.M. and M.T.L conceived the project and wrote the paper. D.D. designed and performed experiments, analyzed data and built the quantitative models. A.G.G. performed covariation analysis for ~350 protein-protein interactions. B.W. helped with library transformations. T.V.L. created the combinatorial antitoxin mutant library. E.N.W. suggested helpful tips on Bayesian modelling. D.S.M. and M.T.L. supervised the project.

## Data and materials availability

Raw sequencing read data will be available upon publication in the Sequencing Reads Archive under accession PRJNA768258.

Analysis code and processed data availability: https://github.com/ddingding/coevolution_paper

