## Supplemental Figures, Table S1, and Methods for "Coevolution of interacting proteins through non-contacting and non-specific mutations"

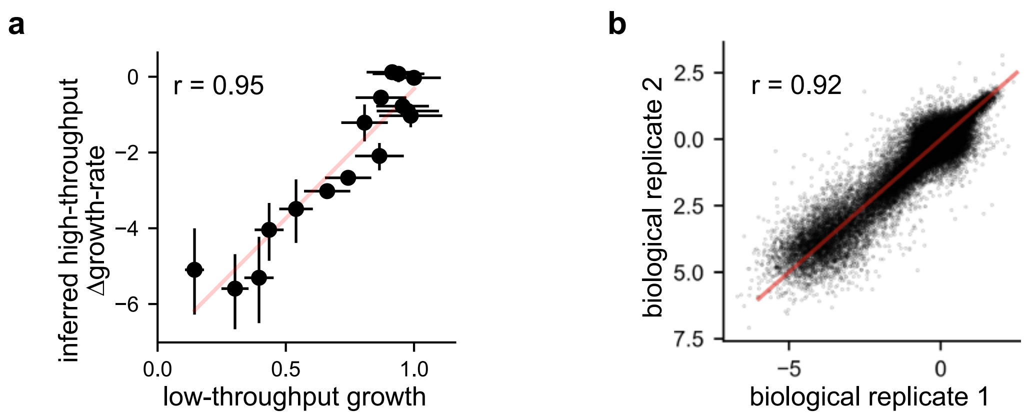


**Extended Data Figure 1**: Orthogonal validation of growth rate inference and reproducibility.

**a**, Comparison of growth rates inferred by high-throughput vs. individual growth measurement. X-axis error bars indicate +/- 2x standard deviation. Y-axis error bars indicate 95% posterior highest density interval. The Pearson correlation coefficient (r) is indicated. Assayed toxin and antitoxin variants (T*+AT*) are: T+AT(G62L), T+AT(W59T), T+AT(F73K), T+AT(K63L), T(V5L)+AT(G62L), T(V5L)+AT(W59T), T(E37D)+AT(G62L), T(G81Y)+AT(G62L), T(P8A)+AT(G62L), T(R52L)+AT(G62L), T(P8N)+AT(W59T), T(V75G)+AT(W59T), T(G94)+AT(W59T), T(W85P)+AT(W59T), T(V5L)+no antitoxin, T(A66F)+no antitoxin.

**b**, Raw log read ratio reproducibility between replicates (+1 pseudocount) for all single and double mutants. The Pearson correlation coefficient (r) is indicated.

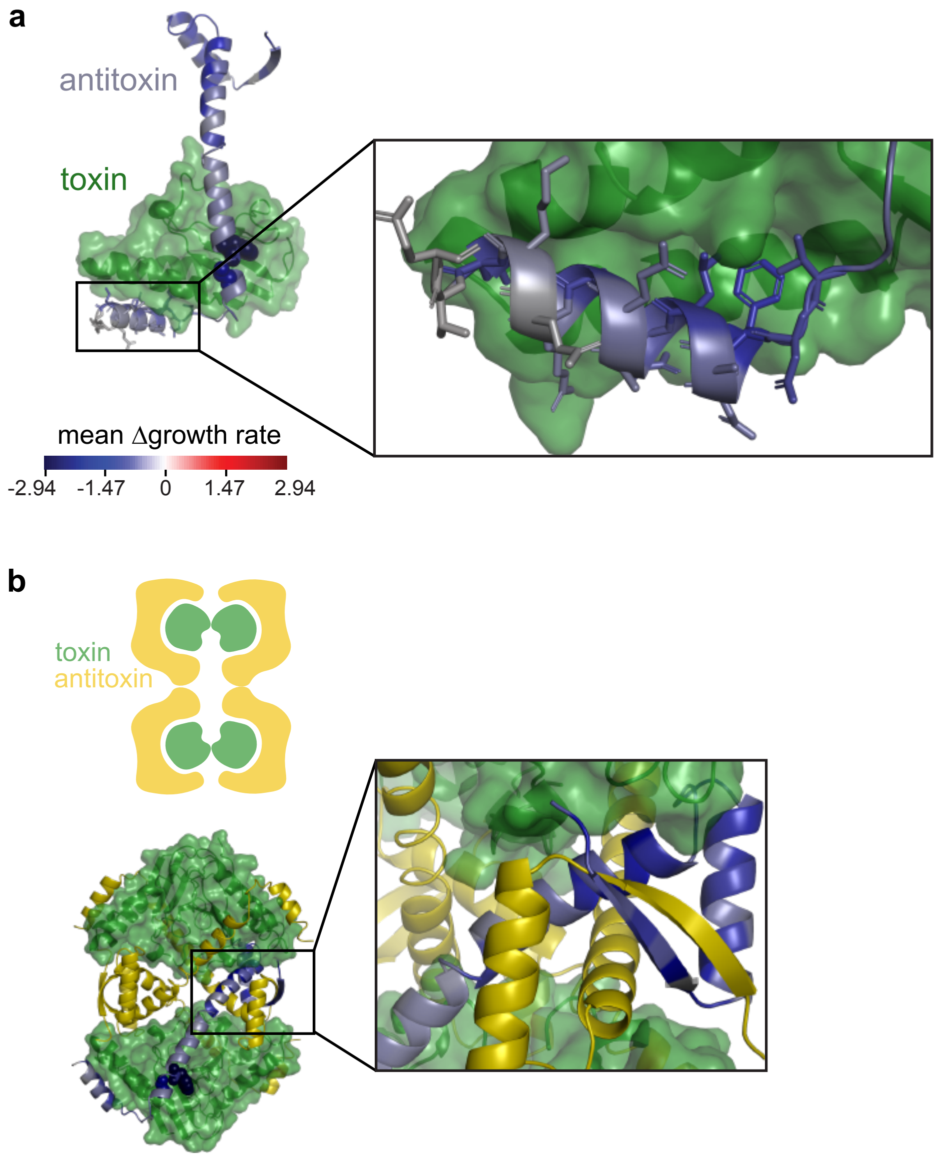


**Extended Data Figure 2.** Avarage position-wise mutation effects in the ParD3 antitoxin reflect requirement of oligmomerization and $\alpha$-helix 3 for toxin neutralization.

**a**, Mean mutation effect of residues in the C-terminal $\alpha$-helix 3 of the ParD3 antitoxin indicates that residues facing the toxin are more susceptible to mutations that disrupt the ParD3-ParE3 interaction, producing negative $\Delta$growth rate values.

**b**, Mean mutation effect in the N-terminal oligomerization region of the antitoxin are susceptible to mutations that disrupt the ParD3-ParE3 interaction. Cartoon illustrates arrangement of ParE3-ParD3 octamer observed in the co-crystal structure (PDB: 5CEG). One of the 4 antitoxin monomers is colored by the mean mutation effect.

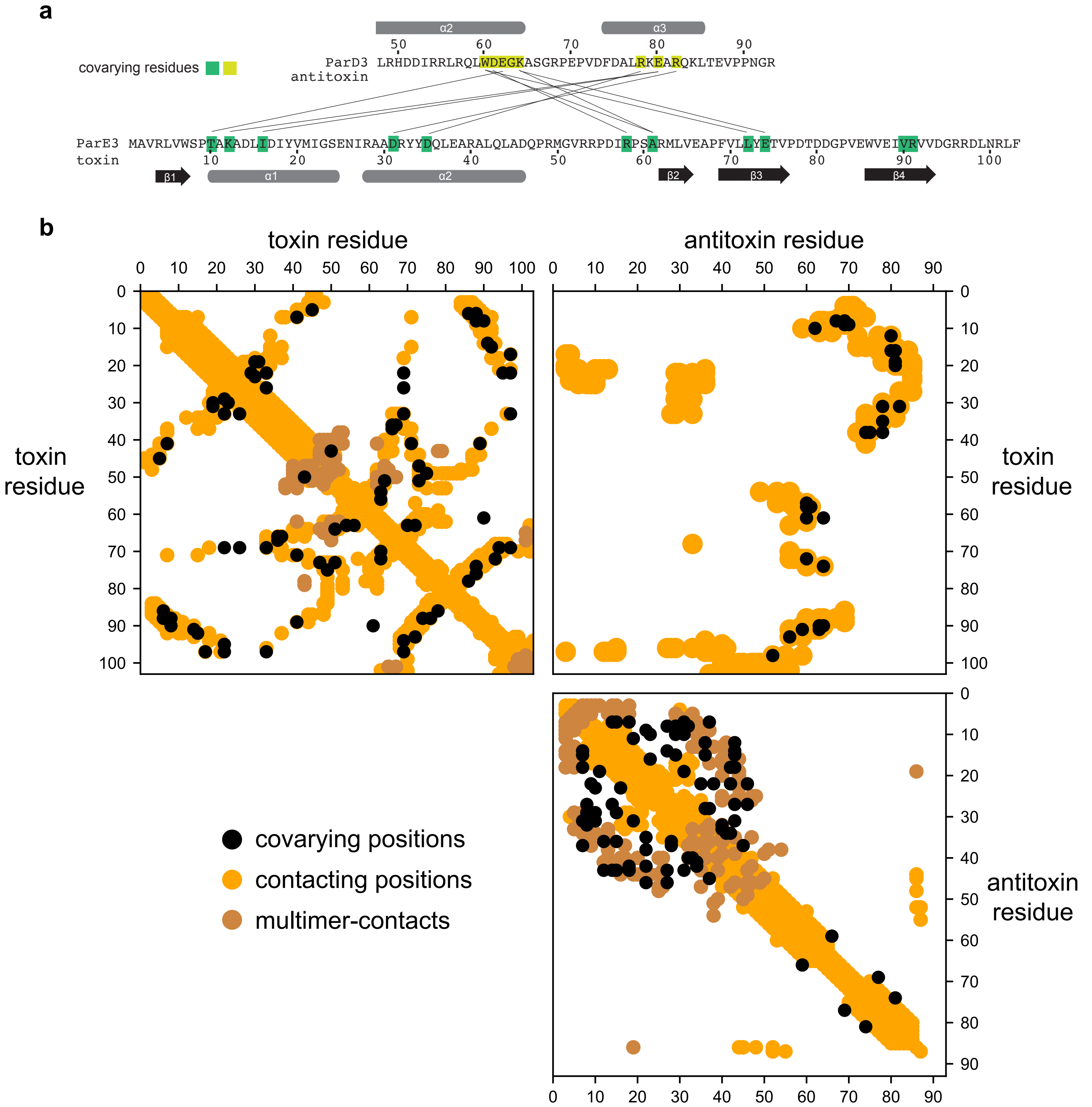


**Extended Data Figure 3:** Covarying residue pairs in natural homologs of ParE3/ParD3 reveal spatially close interface residue pairs.

**a**, Top 10 toxin-antitoxin covarying residue pairs indicated for reference.

**b**, The 90% precision cutoff yields 29 toxin-antitoxin covarying residue pairs (black in upper, right quadrant) of which 28 pairs fall within toxin-antitoxin interface residues that are < 6Å minimum atom distance (ochre dots) in the ParE3-D3 crystal structure (PDB ID: 5CEG).

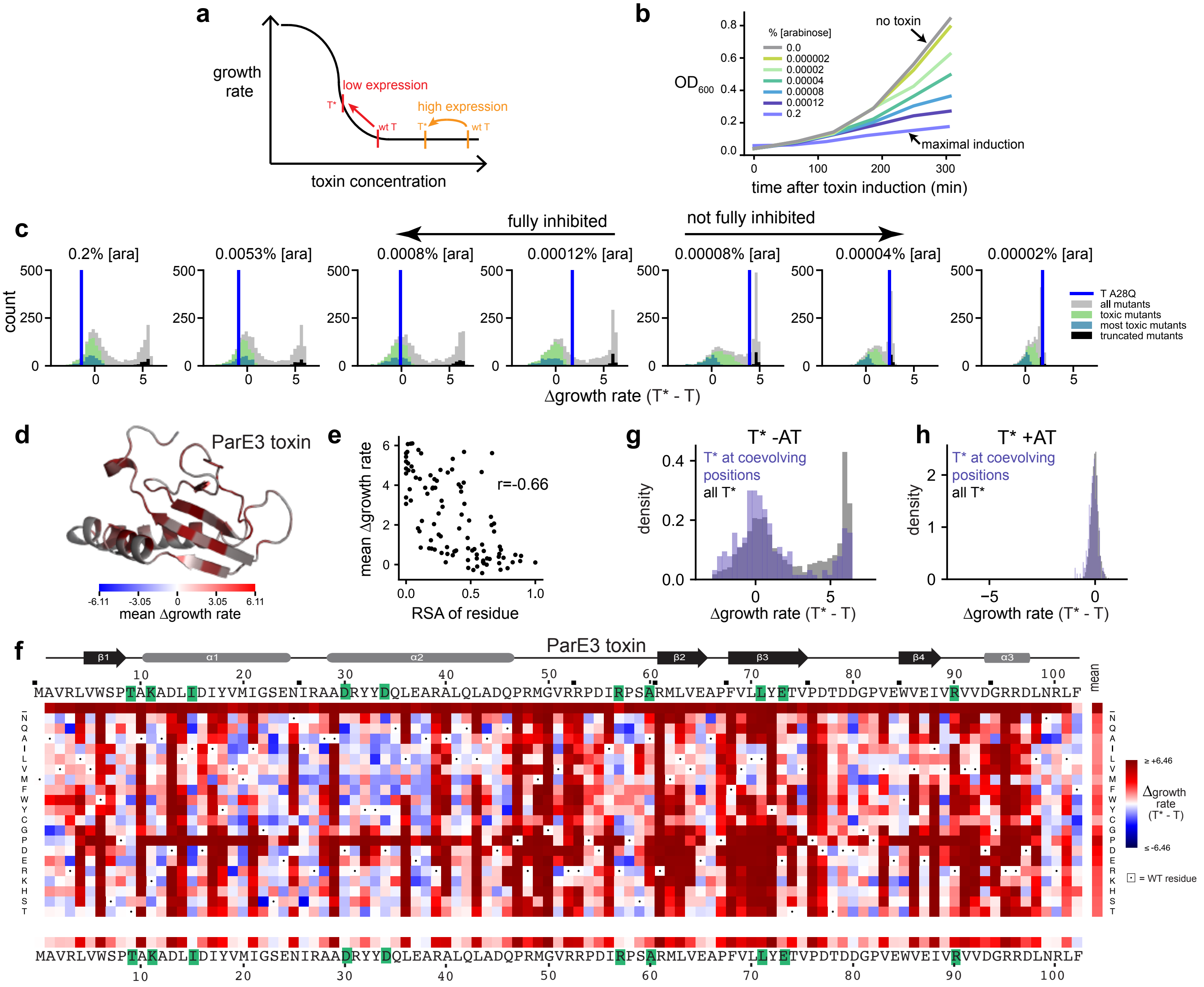


**Extended Data Figure 4:** Sensitive Identification of toxin substitutions which do not disrupt toxicity.

**a**, Schematic illustrating loss of toxicity detection using growth rate measurements in different expression regimes. Measuring toxin variant function at different parts of the growth inhibition curve is necessary to sensitively determine loss of toxicity.

**b**, Growth curves of wild-type toxin in the absence of antitoxin under different toxin induction levels by varying arabinose inducer.

**c**, Distribution of Δgrowth rates(T*-T) for all toxin single substitutions under different arabinose inducer concentrations, with positive Δgrowth rate(T*-T) values indicating loss of toxin function. The set of 'most toxic' toxin subsitutions (n=310) is colored in light blue, the set of 'toxic' substitutions (n=781) is colored in green (see Methods). Other classes of substitutions are indicated. The dynamic range (difference between 0 and the truncated toxin mutants) shrinks, as expected, for lower expression levels that do not fully inhibit growth with the wild-type toxin, and a higher fraction of mutants show loss of toxicity (higher Δ growth rates) under lower expression conditions. The toxin substitution A28Q is highlighted (dark blue) as an example that shows no growth rate difference relative to wild-type toxin at high expression conditions, but is not as toxic as wild-type toxin at lower expression conditions.

**d**, Mean Δgrowth rates(T*-T) of residue positions mapped onto the ParE3 toxin structure. Values shown for 0.00012% [arabinose] inducer.

**e**, The mean Δgrowth rates(T*-T) of a residue are correlated with the relative solvent accessibility of the residue (Pearson r = -0.66). Values shown for 0.00012% [arabinose] inducer.

**f**, The Δgrowth rate(T*-T) values of each substitution at any position along the toxin ParE3. Green highlights the top 10 covarying positions between toxin and antitoxin in natural homologs. Values shown for 0.00012% [arabinose] inducer.

**g,h**, Distribution of Δgrowth rate(T*-T) for all toxin subsitutions (black) or top 10 coevolving residue substitions (purple) in the toxin in absence of antitoxin (**g**) or presence of antitoxin (**h**). Values shown for 0.00012% [arabinose] inducer, and antitoxin is induced with 10 μM IPTG.

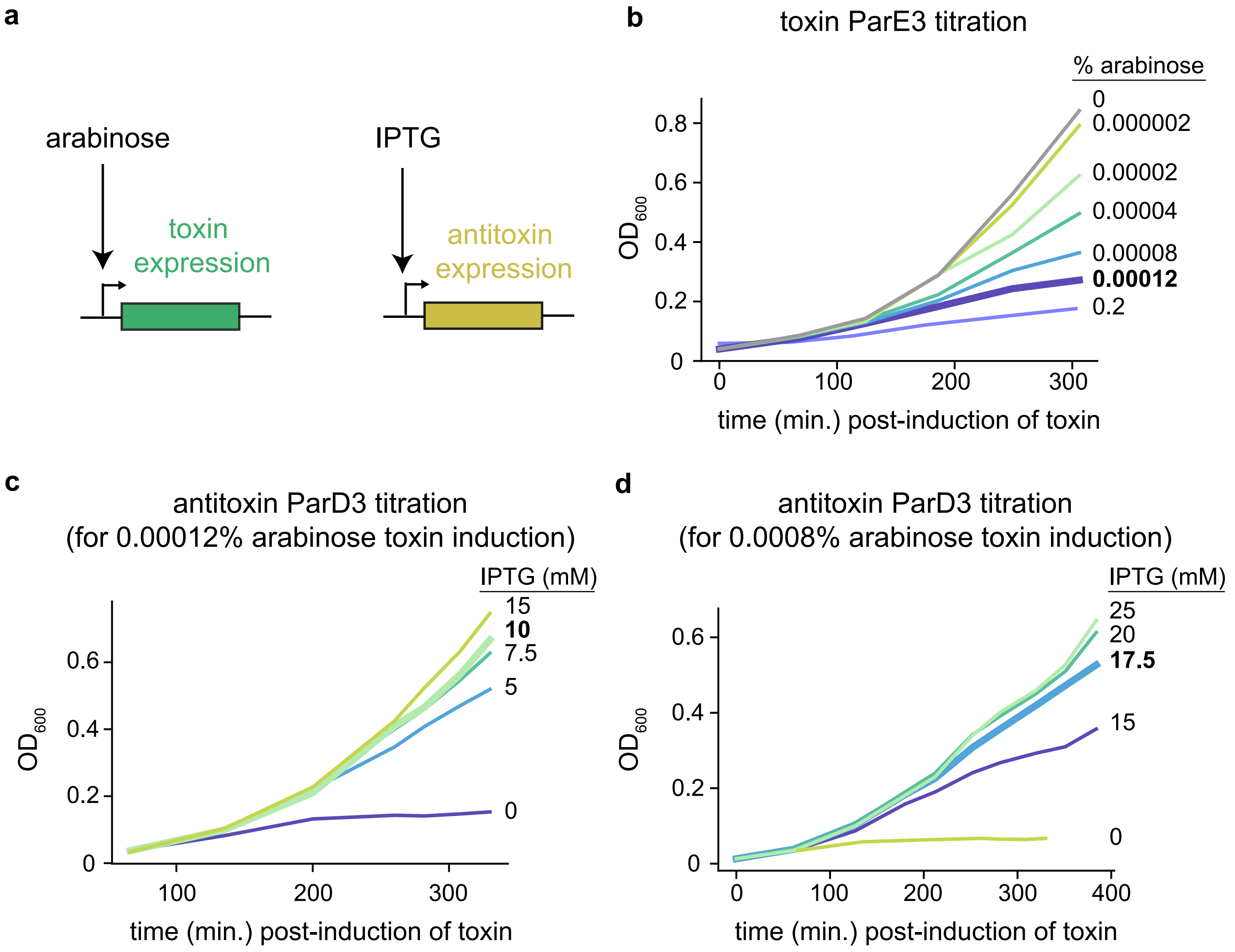


**Extended Data Figure 5:** Determining toxin and antitoxin induction levels for suppressor scanning.

**a**, Cartoon illustration of expresion system. IPTG induces antitoxin, arabinose induces toxin.

**b**, Growth rate of cells harboring wild-type toxin ParE3 without antitoxin at different arabinose induction levels in arabinose titratable *E. coli* strain BW27783.

**c**, Growth rate of cells harboring wild-type toxin-antitoxin ParE3/ParD3 under different antitoxin induction levels modulated with IPTG and 0.00012% arabinose induction.

**d**, Growth rate of cells harboring wild-type toxin-antitoxin ParE3/ParD3 under different antitoxin induction levels modulated with IPTG and 0.0008% arabinose induction.

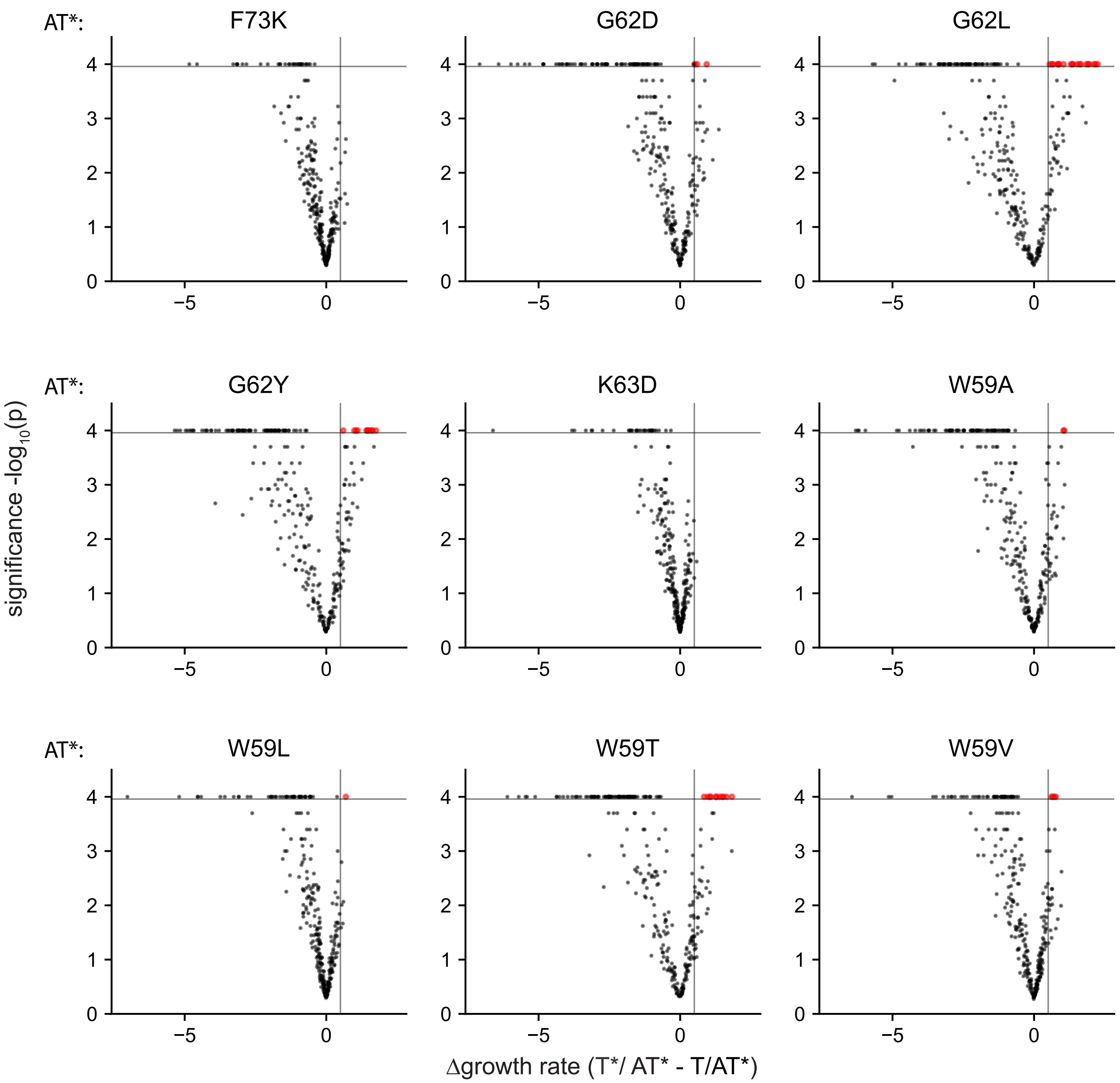


**Extended Data Figure 6:** Volcano plot visualizing significant and substantial beneficial toxin variants in different antitoxin backgrounds.

For each deleterious antitoxin variant background, the mean posterior change in the number of doublings, Δgrowth rate(T*/AT* - T/AT*), of the most toxic toxin mutants are plotted vs. their significance (-log$_{10}$(p(Δgrowth rate<0))) of deviation from the AT* single mutation. This is based on 10,000 discrete samples of the posterior Δgrowth rate(T*/AT* - T/AT*) values inferred from the hierarchical Bayesian inference model (see Methods). Vertical line: +0.5 Δgrowth rate, horizontal line: p(Δgrowth rate>0) = 0.0001. Red indicates significant and substantial beneficial toxin substitution. Experiments performed under 'high antitoxin' expression conditions.

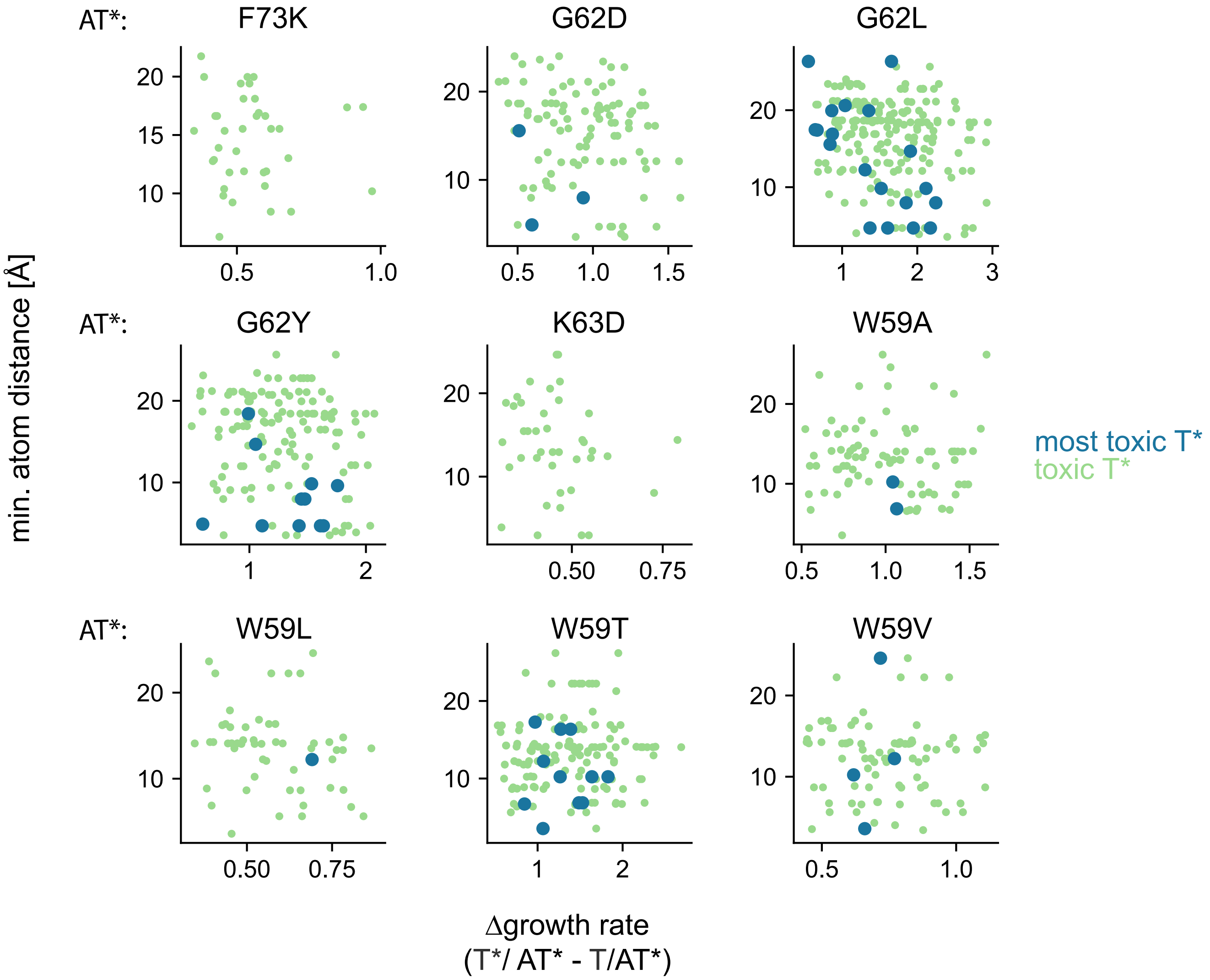


**Extended Data Figure 7:** Distance of rescuing toxin substitutions for each deleterious antitoxin substitution under 'high antitoxin' expression conditions.

The minimum atom distance from a given deleterious antitoxin residue to each beneficial toxin is plotted vs. Δgrowth rate(T*/AT* - T/AT*). Experiments performed under 'high antitoxin' expression conditions.

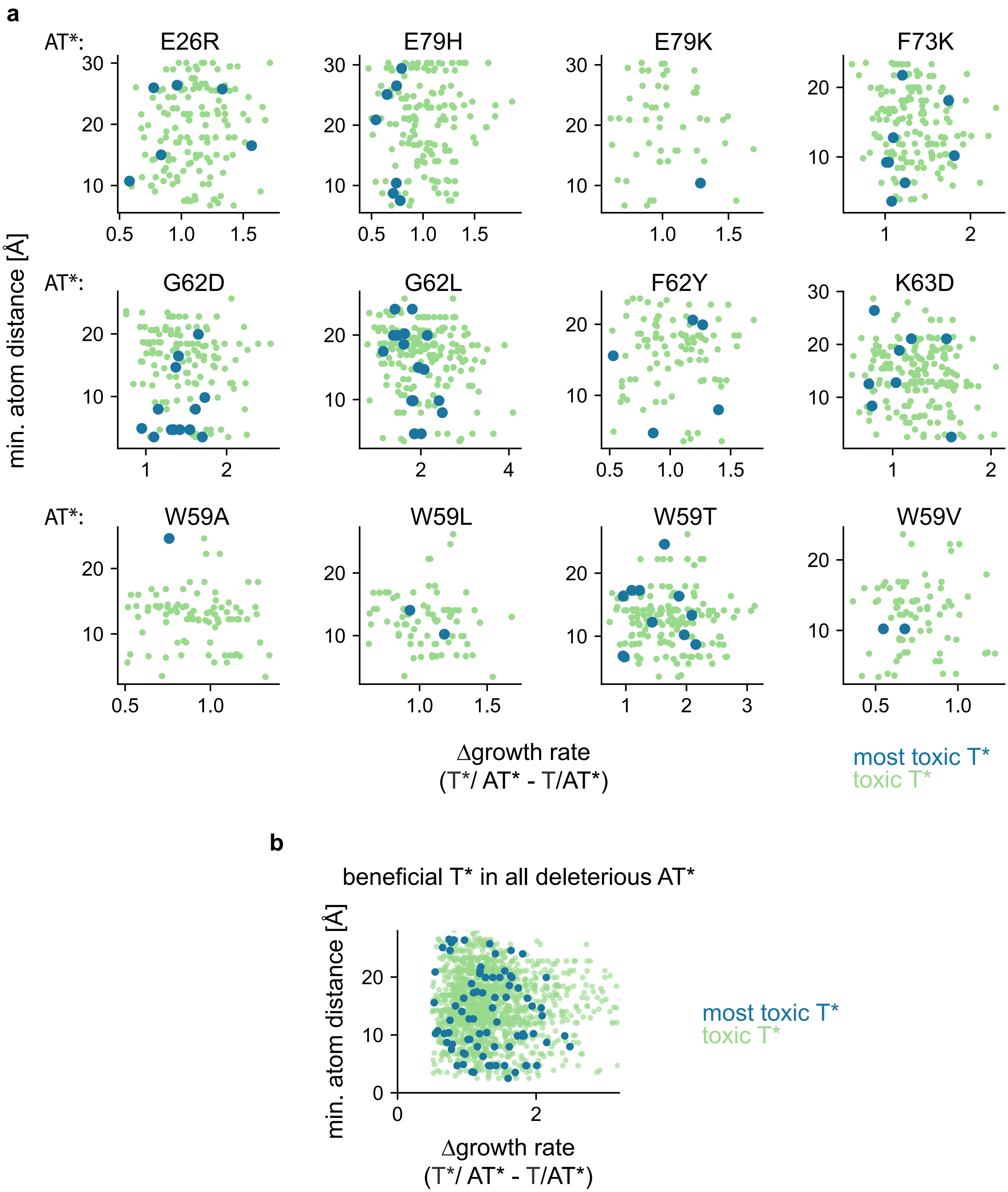


**Extended Data Figure 8:** Distance of rescuing toxin substitutions for each deleterious antitoxin substitution under 'low antitoxin' expression conditions.

**a**, The minimum atom distance from a given deleterious antitoxin residue to each beneficial toxin is plotted vs. Δgrowth rate(T*/AT* - T/AT*). Experiments performed under 'low antitoxin' expression conditions.

**b**, Distance vs. Δgrowth rate(T*/AT* - T/AT*) of beneficial toxin variants for all deleterious antitoxin variant backgrounds.

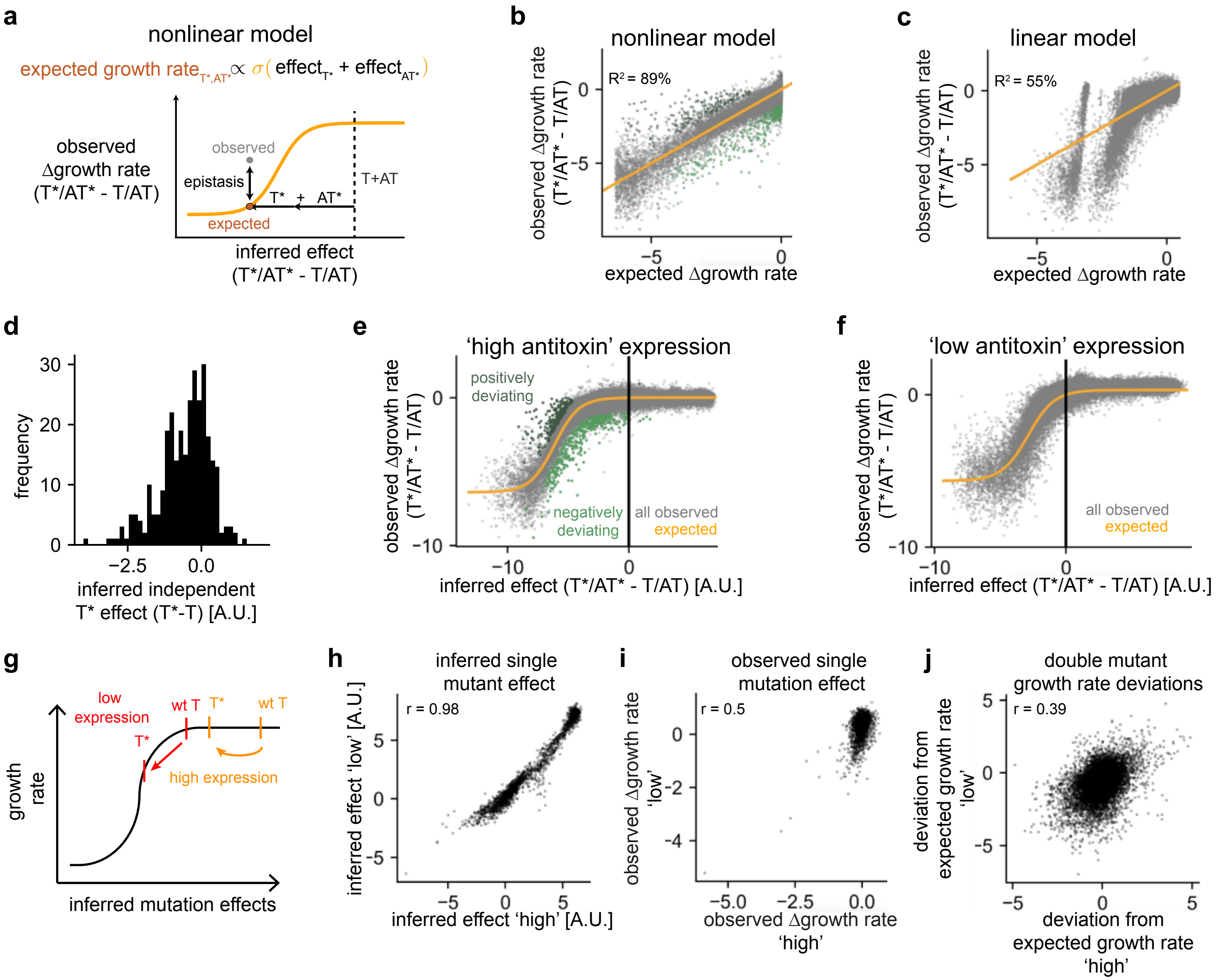


**Extended Data Figure 9:** A non-specific, non-linear model can explain most of the observed single and double mutant growth rates.

**a**, Schematic of nonlinear, non-specific model: double mutant expected growth rates (brown) are based on the independent (non-specific) sum of underlying toxin and antitoxin mutant effects, passed through a sigmoid function (yellow).

**b,c**, Residuals for non-linear, non-specific model (**b**) or linear non-specific model of the same structure without a non-linearity (**c**) showing unbiased residuals for the nonlinear model, but a complete misfit of the linear model. Model built using 'high antitoxin' expression levels. Explained variance ($R^2$) is indicated. Significant and substantially positively(dark green) or negatively (green) deviating mutations are shown in (**b**) (see Methods).

**d**, Inferred independent toxin single substitution effects among the set of most toxic toxin mutants demonstrating a tail of independently beneficial toxin variants. Experiment performed under 'high antitoxin' expression levels.

**e,f**, Nonlinear independent model fit to growth rates measured under 'high antitoxin' (**e**) or 'low antitoxin' (**f**) expression conditions. The wild-type toxin -antitoxin pair is inferred to be differently close to the sigmoid 'cliff' between expression conditions.

**g**, Cartoon illustrating different detection of single mutant effects depending on expression conditions.

**h-j**, Correlation of inferred single mutant effects (**h**), observed single mutant Δgrowth rate(T*/AT* - T/AT) effects (**i**), and double mutant deviations of observed from expected growth rates (**j**) from separate inference under 'high antitoxin' (x-axis) or 'low antitoxin' (y-axis) expression conditions.

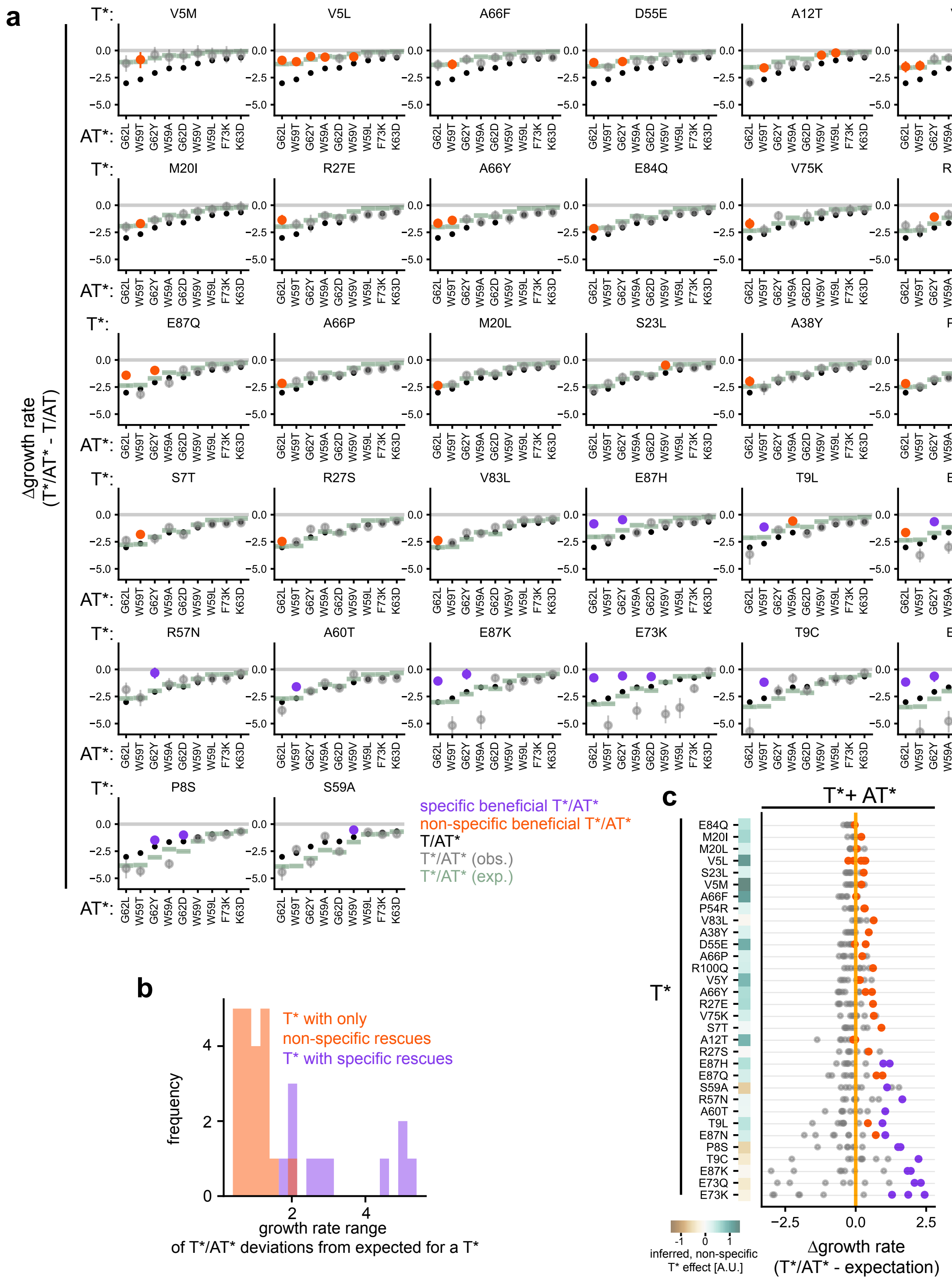


**Extended Data Figure 10:** Deviation of observed from expected double mutant growth rates reveals toxin variants with specific or with only non-specific beneficial effects.

**a**, For each beneficial toxin mutation (indicated above each plot) combined with each antitoxin variant indicated on the x-axis, the plot shows the growth-rate relative to the wild-type toxin-antitoxin pair (Δgrowth rate(T*/AT* - T/AT)). Grey dots represents T*/AT*, error bars indicate 95% posterior highest density interval. The Δgrowth rate for each antitoxin mutant combined with wild-type toxin (T/AT*) is shown (black dots) along with the Δgrowth rate for T*/AT* expected under the non-specific, nonlinear model (green dots).

**b**, Distribution of the range of deviations from expected growth rates for toxin substitutions across deleterious antitoxin mutants (Δgrowth rate(AT*/T*)-expected growth rate) revealig separation of toxin variants containing specific (purple) or only non-specific rescues (orange).

**c**, Deviation of the observed (dots) from the expected double mutant growth rates (orange line) highlights classification of specific and non-specific toxin variants. Beneficial toxin subsitutions (rows, n=32) ordered by their range of growth rate deviations across deleterious antitoxin variants as in panel b.

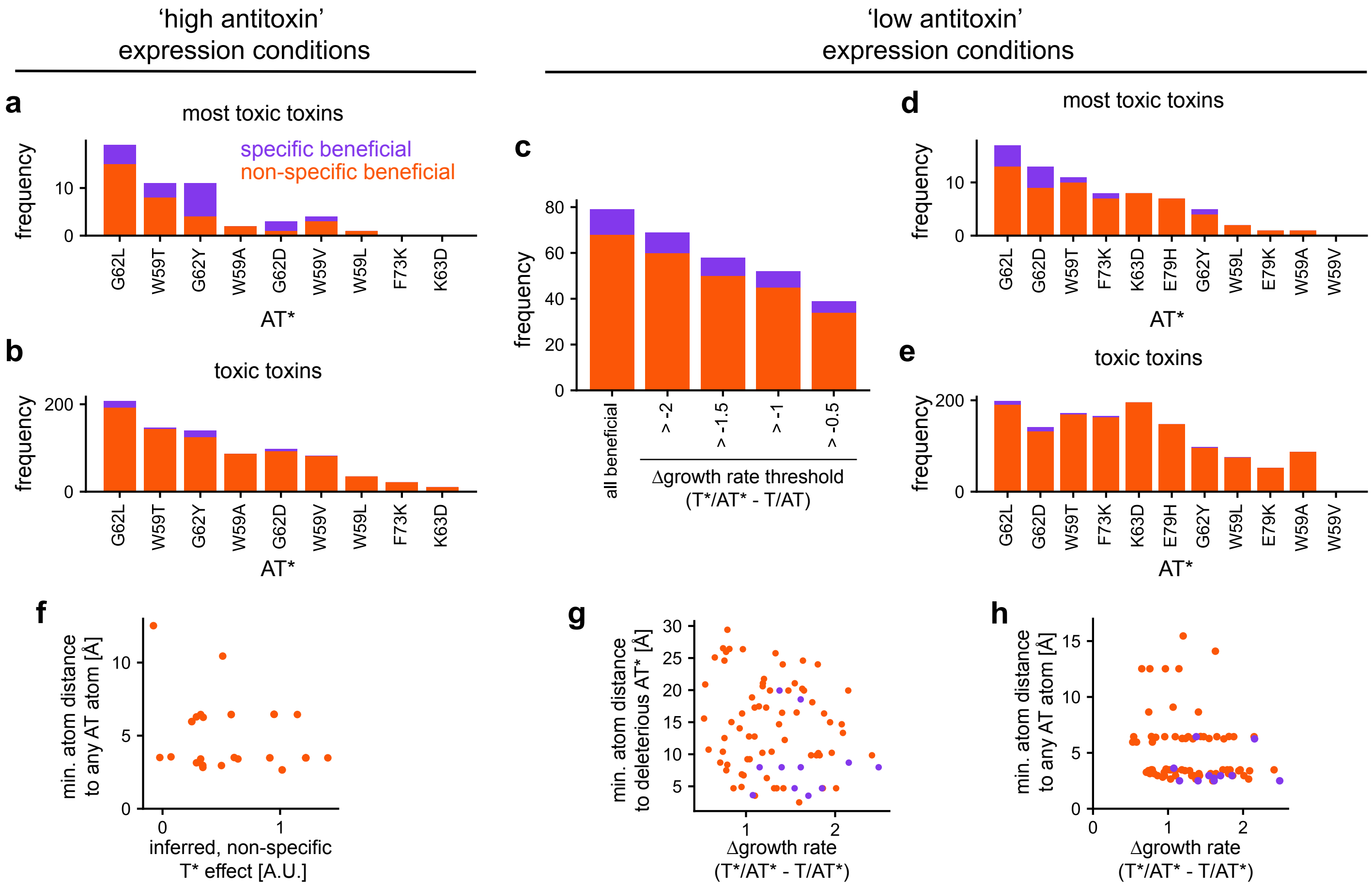


**Extended Data Figure 11:** Number and spatial distribution of specific and non-specific toxin variants under different expression conditions.

**a,b**, Specific vs. non-specific enabling toxin variants under 'high' antitoxin expression for all enabling toxin variants grouped by deleterious antitoxin for the more stringent set of 310 'most toxic' toxins (**a**) and less stringent set of 781 'toxic' toxins (**b**). Orange and purple indicate mutant pairs involving non-specific and specific, respectively, rescuing mutations in the toxin.

**c-e**, Specific vs. non-specific enabling toxin variants under 'low' antitoxin expression for all enabling toxins, or at different absolute growth rate cutoffs relative to the wild-type toxin/antitoxin growth rate (**c**), grouped by antitoxin for the set of 310 'most toxic' toxin mutants (**d**) or 781 'toxic' toxins (**e**). Specific and non-specific pairs color coded as in A-B.

**f**, Inferred non-specific toxin variant effect vs. minimum atom distance to any antitoxin atom for 21 non-specifically rescuing toxin variants (orange). 8 non-specific toxin subsitutions (from 6 different positions) are >6 Å minimum atom distance from the antitoxin, and 13 non-specific toxin substitutions (from 9 different positions) are <6 Å minimum atom distance to any antitoxin atom.

**g,h**, For specific and non-specific beneficial toxin mutants, the change in growth rate in a deleterious antitoxin mutant background, Δgrowth rate (T*/AT* - T/AT*), is plotted vs. minimum atom distance to the deleterious antitoxin mutation it rescues (**g**) or any antitoxin atom (**h**) in the 'low antitoxin' expression condition.

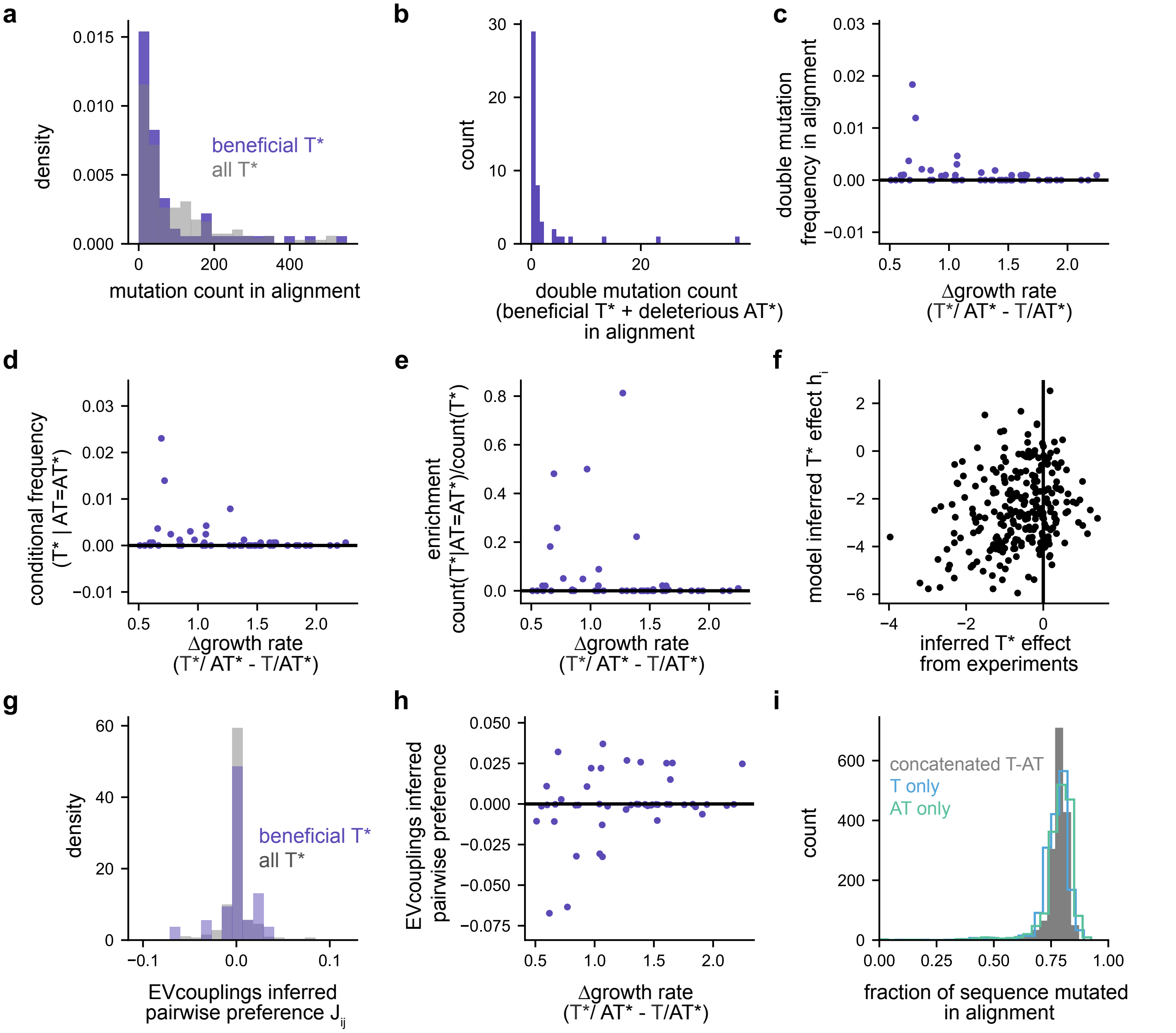


**Extened Data Figure 12:** Natural sequence statistics and EVcouplings model parameters are not predictive of beneficial toxin substitution effects.

**a**, Distribution of number of specific and non-specific beneficial toxin substitutions (purple) vs. all possible toxin variants (grey) observed in natural sequences.

**b**, Frequency distribution of beneficial toxin and deleterious antitoxin mutant pairs in natural sequences, with 29/51 pairs never observed.

**c-e**, Effect size of toxin variant rescue vs. frequency of variant pair in natural sequences (**c**), conditional frequency of toxin variant given natural sequences containing the particular deleterious antitoxin substitution (**d**), or enrichment of beneficial toxin variant in natural sequences containing the deleterious antitoxin substitution (**e**).

**f-h**, EVcouplings model inferred sitewise toxin mutant preferences ($h_i$) vs. toxin mutant effect inferred in suppressor scan (**f**), EVcouplings pairwise variant preference ($J_{ij}$) distribution for variant pairs consisting of a beneficial toxin and deleterious antitoxin mutant pair compared to random variant pairs (**g**), or EVcouplings pairwise T*/AT* variant preference ($J_{ij}$) vs. effect size of beneficial toxin mutation effect in a deleterious antitoxin variant background (**h**).

**i**, Distribution of natural sequence identity fractions across the alignment. Different histograms illustrate fraction mutated for homologs containing the full concatenated toxin and antitoxin (grey), the toxin homologs only (blue), or the antitoxin homologs only (turqouise).

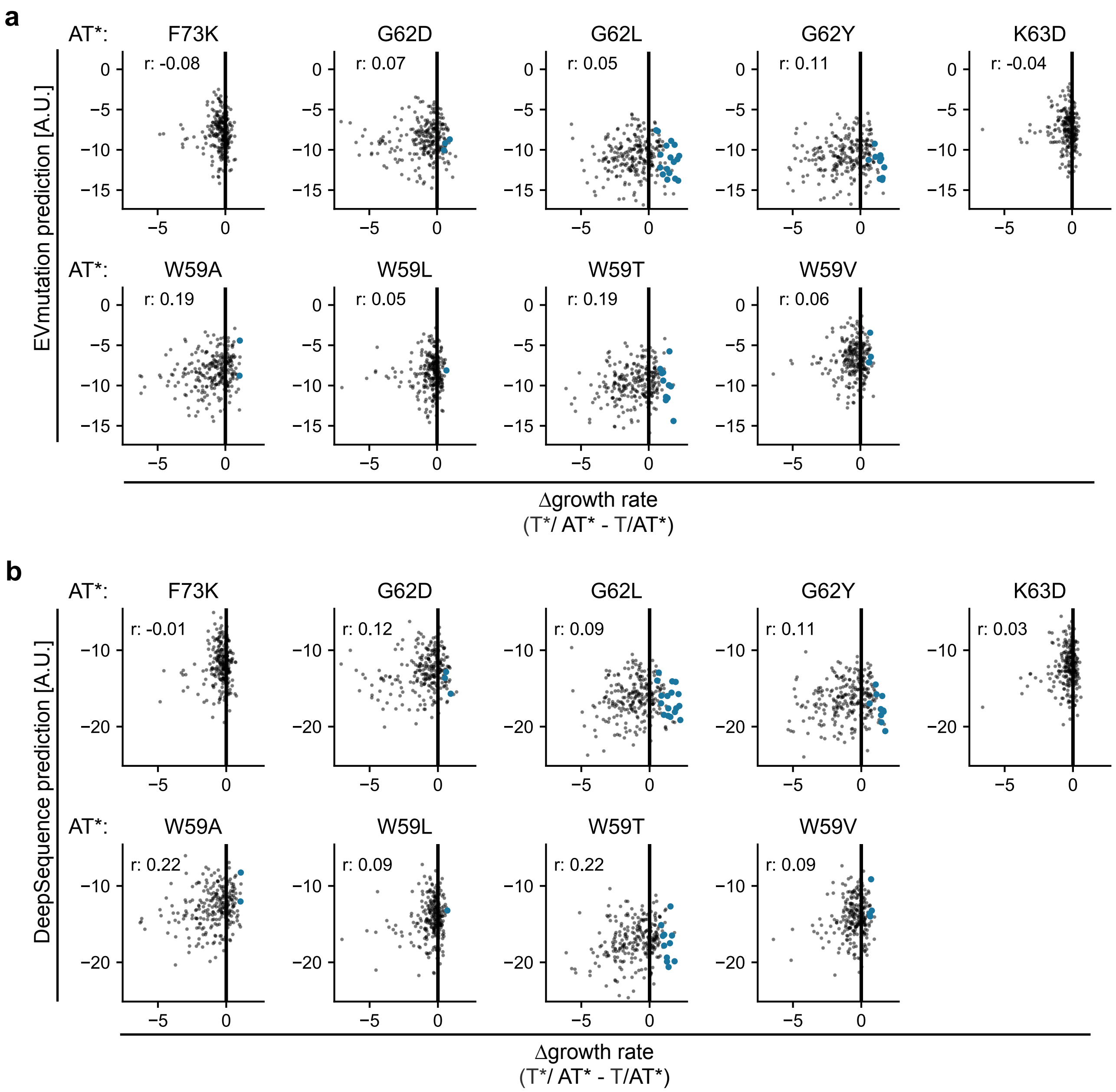


**Extended Data Figure 13:** Models trained on a sequence alignment cannot predict beneficial toxin variants in any deleterious antitoxin variant background.

**a,b**, Scatterplot of observed beneficial toxin effect in deleterious antitoxin mutant backgrounds (AT*), vs EVmutation (**a**) or DeepSequence (variational autoencoder) mutation effect predictions (**b**). Pearson correlation (r) is indicated.

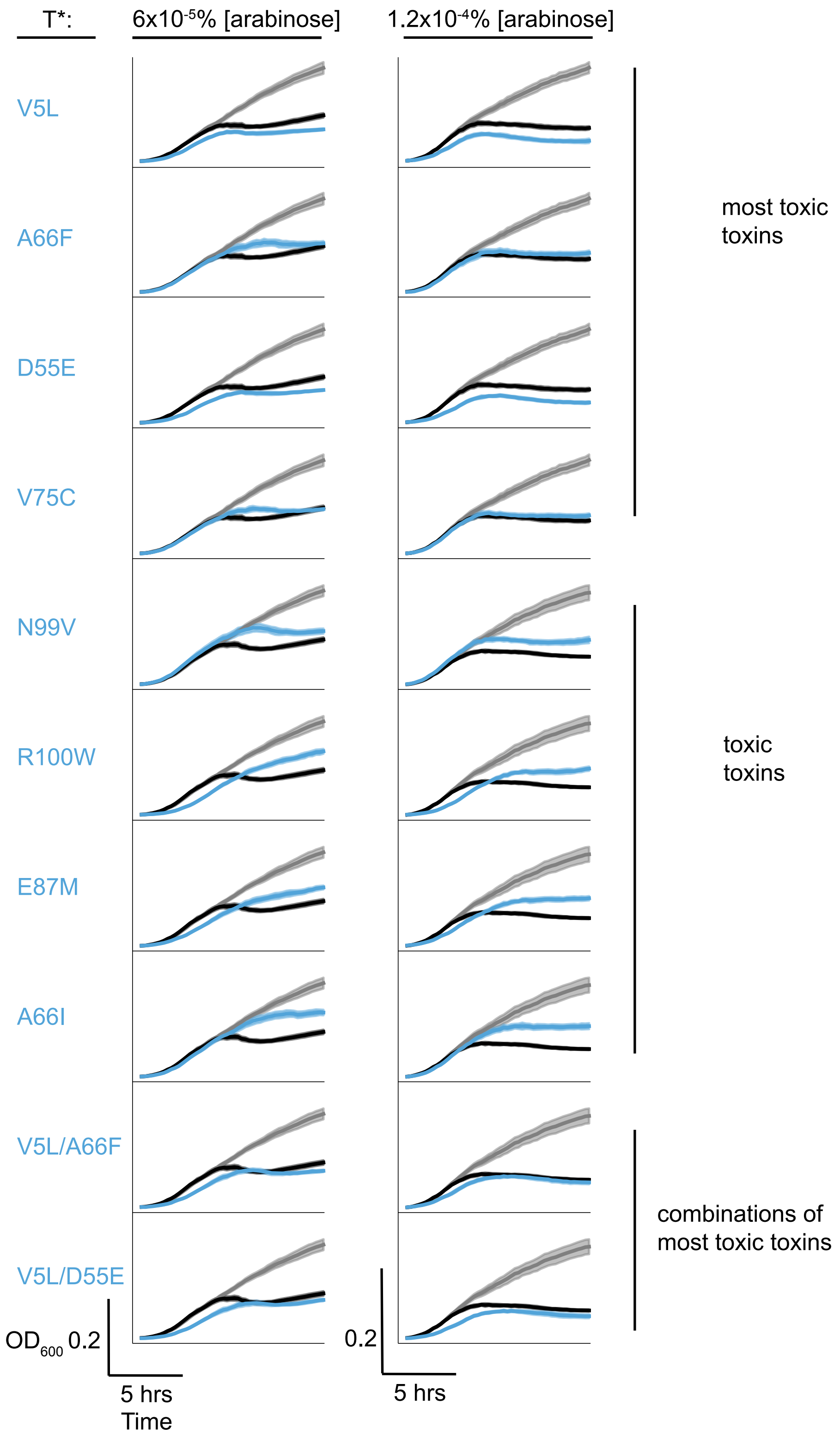


**Extended Data Figure 14:** Non-specific suppressor toxin ParE3 variants are as or almost as toxic as wild-type ParE3.

Growth rates of ParE3 non-specific suppresor toxin variants (blue) compared to wild-type toxin ParE3 without antitoxin (black) and wild type toxin and antitoxin (grey) under fully inhibitory toxin expression conditions (0.00012% [arabinose]) or half-maximal inhibitory expression conditions (0.00006% [arabinose]). Shaded regions show standard deviation of the replicates (n=10-12).

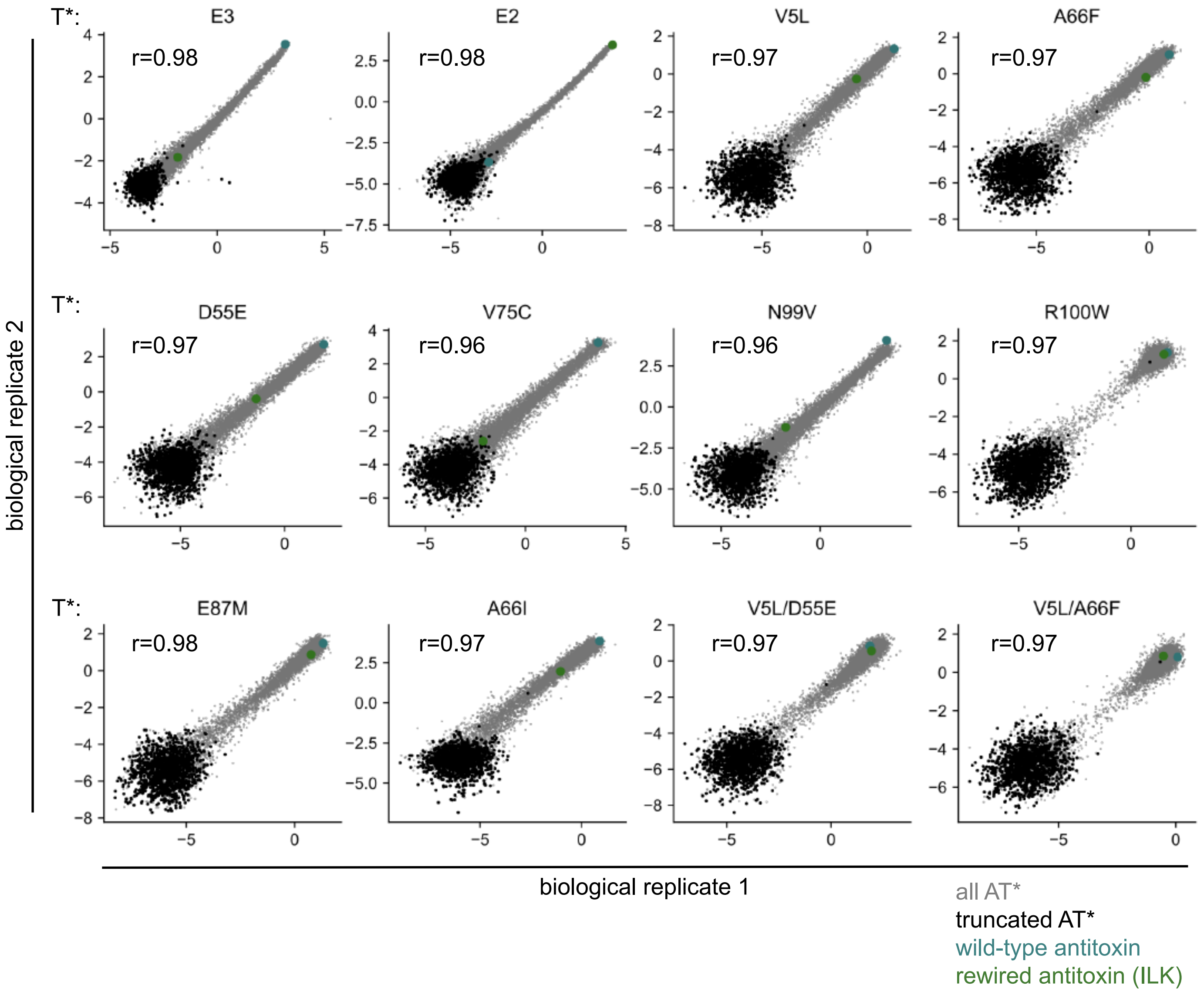


**Extended Data Figure 15:** Reproducibility of antitoxin combinatorial variant log read ratios.

Raw log read ratio reproducibilities between biological replicates (+1 pseudocount) for the combinatorial antitoxin library (8000 amino acid variants) in different toxin mutant backgrounds. Specific classes of antitoxin mutants, and Pearson correlation coefficients (r) are indicated.

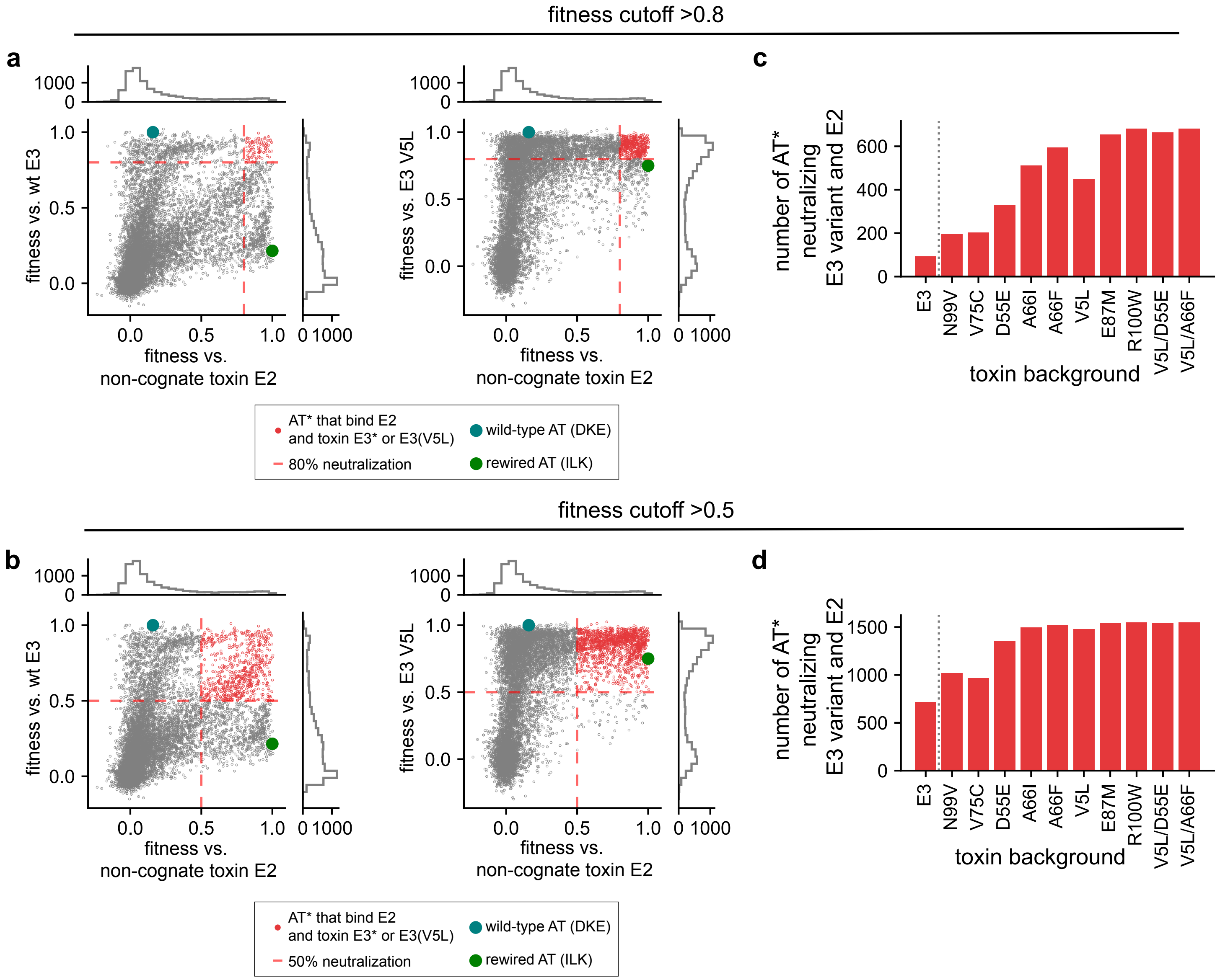


**Extended Data Figure 16:** Non-specific suppressors in toxin ParE3 enable binding to ParD3 variants that also bind non-cognate ParE2.

**a,b**, Scatterplots showing fitness values for each of 8,000 antitoxin variants screened against the wild-type toxin ParE3 and the non-cognate toxin ParE2, or against toxin ParE3(V5L) and non-cognate ParE2 at a fitness cutoff of 80% (**a**) or 50% (**b**) maximal neutralization. Various classes of AT* are color-coded as shown beneath, including the AT* variants that promiscuously bind both toxins (red).

**c,d**, Number of promiscuous antitoxin variants that neutralize both the toxin indicated on the x-axis as well as non-cognate ParE2 at a fitness cutoff of 80% (**c**) or 50% (**d**).

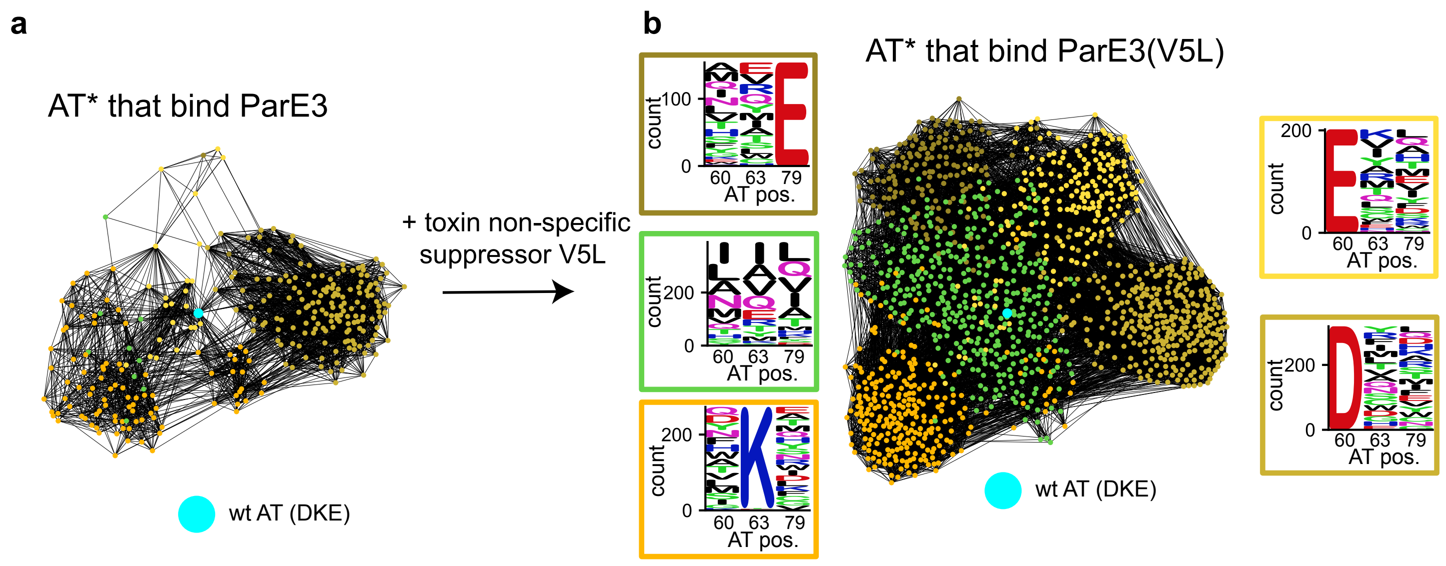


**Extended Data Figure 17:** Accessible antitoxin mutants in the presence and absence of the toxin non-specific suppressor V5L.

**a,b**, Force-directed graph of antitoxin variants(nodes) with >90% of wild-type antitoxin neutralization when expressing the wild-type toxin ParE3 (A) or the variant ParE3(V5L). Nodes are antitoxin variants, edges represent single mutational steps. Sequence logos of 5 Louivain clusters of antitoxin variants are indicated.

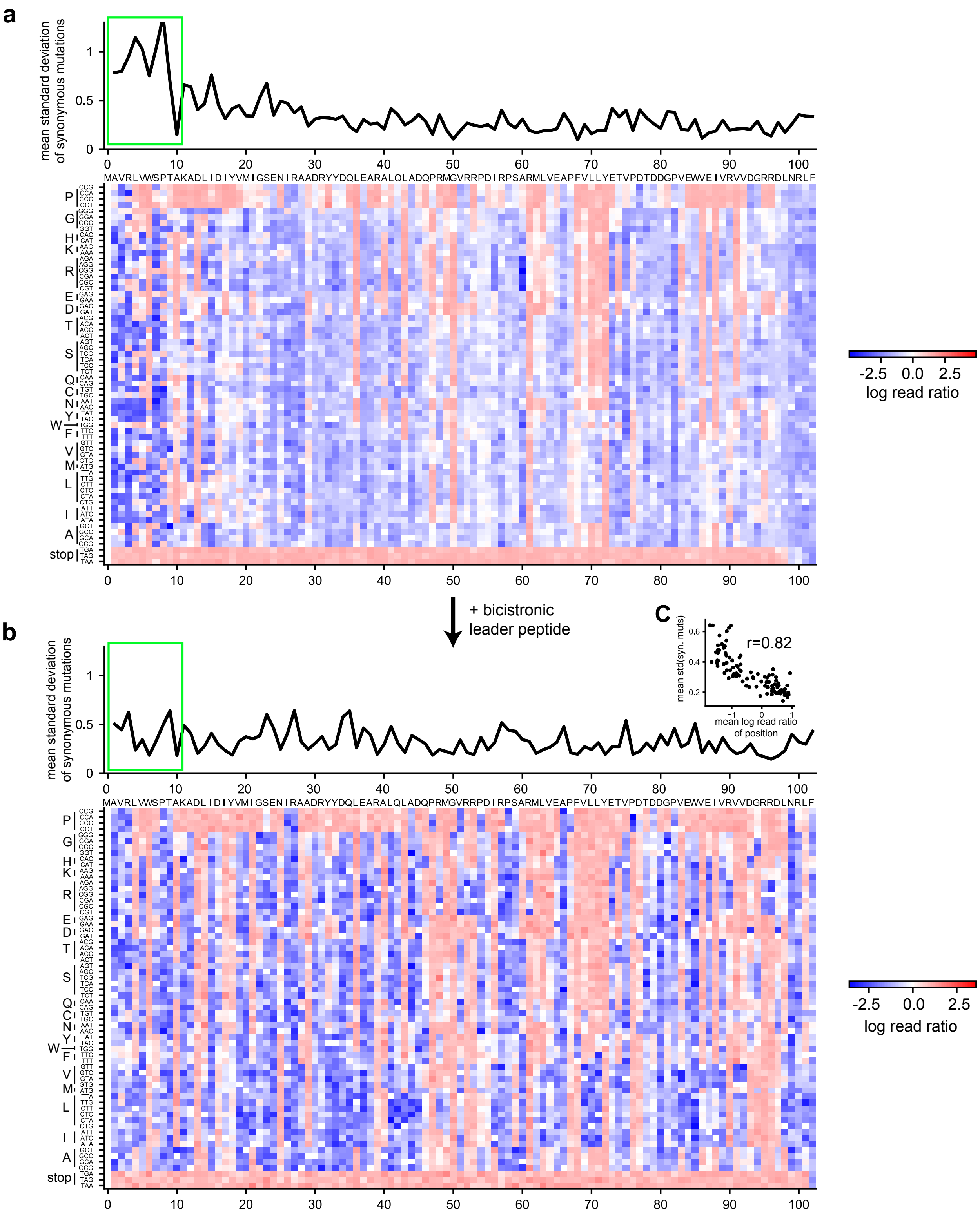


**Extended Data Figure 18:** An upstream bicistron leader peptide removes N-terminal synonymous mutant growth rate variation.

**a**,**b**, Observed log read ratios (+1 pseudocount) for toxin single codon library (no antitoxin) without (**a**) or with (**b**) a bicistronic leader peptide. Top of each panel shows the average standard deviation of synonymous mutant log read ratios along the gene, highlighting the N-terminus (green box). Heat maps below show the log read ratios for each codon at each position in the toxin.

**c**, The standard deviation of synonymous toxin single codon variant growth rates in the presence of the bicistronic leader peptide is well explained by the mean log read ratio of that position (Spearman r = 0.82). This can be rationalized since positions with mostly toxicity preserving mutations show higher growth rate variability due to sampling noise at the post-selection timepoint with few observed reads.

**a**

observed reads → amino acid substitution growth rate

$reads_{codon,\ before} \sim Poisson(\lambda_{codon,\ before})$

$reads_{codon,\ after} \sim Poisson(\lambda_{codon,\ after})$

$\lambda_{codon,\ after} = ratio_{codon} \times \lambda_{codon,\ before}$

$\log(ratio_{codon}) \sim Normal(w_{amino\ acid}, \sigma^2_{synonymous})$

**b**

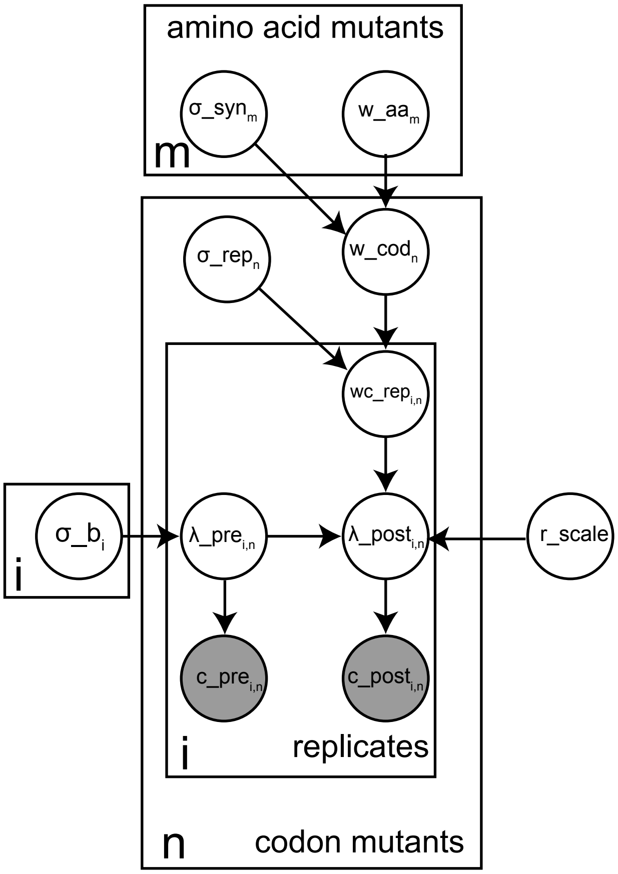


**c**

**Variables**

| | |
|---|---|
| i | replicate index |
| m | amino acid index |
| n | codon index |
| $w\_aa_m$ | amino acid mutant growth rate |
| $\sigma\_syn_m$ | synonyous codon variance |
| $w\_cod_n$ | codon mutant growth rate |
| $wc\_rep_{i,n}$ | replicate codon mutant growth rate |
| $\sigma\_rep_n$ | replicate variance |
| $c\_pre_{i,n}$ | observed codon pre-selection counts |
| $c\_post_{i,n}$ | observed codon post-selection counts |
| $\lambda\_pre_{i,n}$ | pre-selection rate |
| $\lambda\_post_{i,n}$ | post-selection rate |
| $\sigma\_b_i$ | pre-selection rate variance |
| r_scale | replicate scale |

**Likelihood function**

| | |
|---|---|
| $c\_pre_{i,n}$ | $\sim Poisson(\lambda\_pre_{i,n})$ |
| $c\_post_{i,n}$ | $\sim Poisson(\lambda\_post_{i,n})$ |
| $\lambda\_pre_{i,n}$ | $\sim lognormal(0,\sigma\_b_i)$ |
| $\lambda\_post_{i,n}$ | $= wc\_rep_{i,n} \bullet \lambda\_pre_{i,n} \bullet r\_scale$ |
| $wc\_rep_{i,n}$ | $\sim Normal(w\_cod_{n,}\ \sigma\_rep_n)$ |
| $w\_cod_n$ | $\sim Normal(w\_aa_{m,}\ \sigma_{syn})$ |

**Priors**

| | |
|---|---|
| $w\_aa_m$ | ~ Normal(-2,5) |
| $\sigma\_syn_m$ | ~ Exponential(8) |
| $\sigma\_rep_n$ | ~ Exponential(3) |
| $\sigma\_b_i$ | $\sim Uniform(0,\infty)$ |
| r_scale | $\sim Uniform(0,\infty)$ |

**Extended Data Figure 19:** Bayesian hierarchical model.

**a**, Simplified description of the bayesian hierarchical model. Pre- and post-selection reads for each codon are drawn from a Poisson distribution. The log-ratios of these Poisson parameters are not fixed between synonymous codons but are instead drawn from a normal distribution, whose mean forms the amino acid mutant growth rate of interest. This model allows for different synonymous codons to inform each other as well as the amino acid mutant growth rate without being completely fixed.

**b**, Full plate diagram description of the hierarchical Bayesian model capturing both replicates. Replicate index i takes values 1 or 2, amino acid index m takes on values ranging from 1-2040 (20*102) for the toxin or 1-1840 (92 * 20) for the antitoxin, codon index n takes on values ranging from 1-6426 (63*102) for the toxin or 1-5796 (63*92) for the antitoxin. Circles indicate random variables, grey circles represent observed random variables.

**c**, Description of variables, likelihood function and priors used. The likelihood function incorporates maximum entropy distributions for the observed variables, and the priors incorporate computationally tractable, vague priors for the amino acid substitution growth rates. The relative priors on the standard deviation of replicate $\sigma\_rep_n$ vs. synonymous variant $\sigma\_syn_m$ reflect our prior belief that replicate experiment noise is larger than synonymous mutant noise. $\sigma\_b_i$ and r_scale have improper priors.

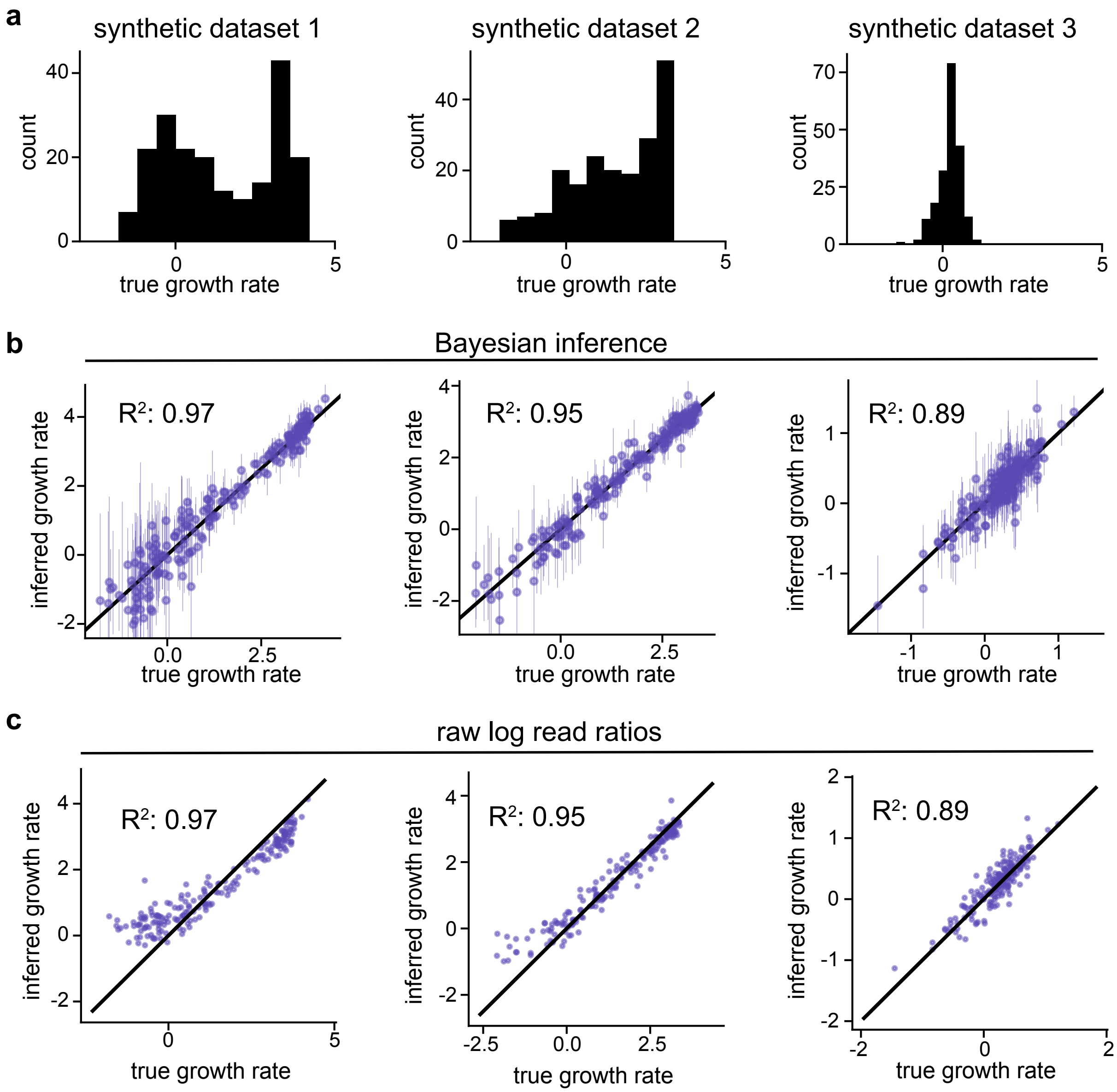


**d**

| | *Dataset 1* | *Dataset 2* | *Dataset 3* |
|---|---|---|---|
| *95% HDI* | *93%* | *96%* | *94%* |
| *90% HDI* | *90%* | *90%* | *91%* |
| *80% HDI* | *75%* | *79%* | *81%* |
| *50% HDI* | *44%* | *44%* | *50%* |

**Extended Data Figure 20:** Validation of Bayesian growth rate inference on synthetic datasets.

**a**, Three different true synthetic growth rate distributions used for simulating pre- and post-selection codon variant read count data. Synthetic growth rate distributions were chosen from observed toxin single mutant growth rate distributions in 3 different antitoxin backgrounds, spanning the range of distributions observed.

**b,c**, Inferred growth rates using the Bayesian hierarchical model (**b**) show less bias and incorporate uncertainty estimates compared to mean log read ratio summary of pre-and post-selection read counts (+1 pseudocount) (**c**). Error bars in panel b reflect the 95% highest density posterior intervals.

**d**, Model uncertainties accurately reflect deviations of inferred true growth rates. Percentage of true synthetic amino acid growth rates falling into a certain highest density interval among all 2040 simulated toxin amino acid variants.

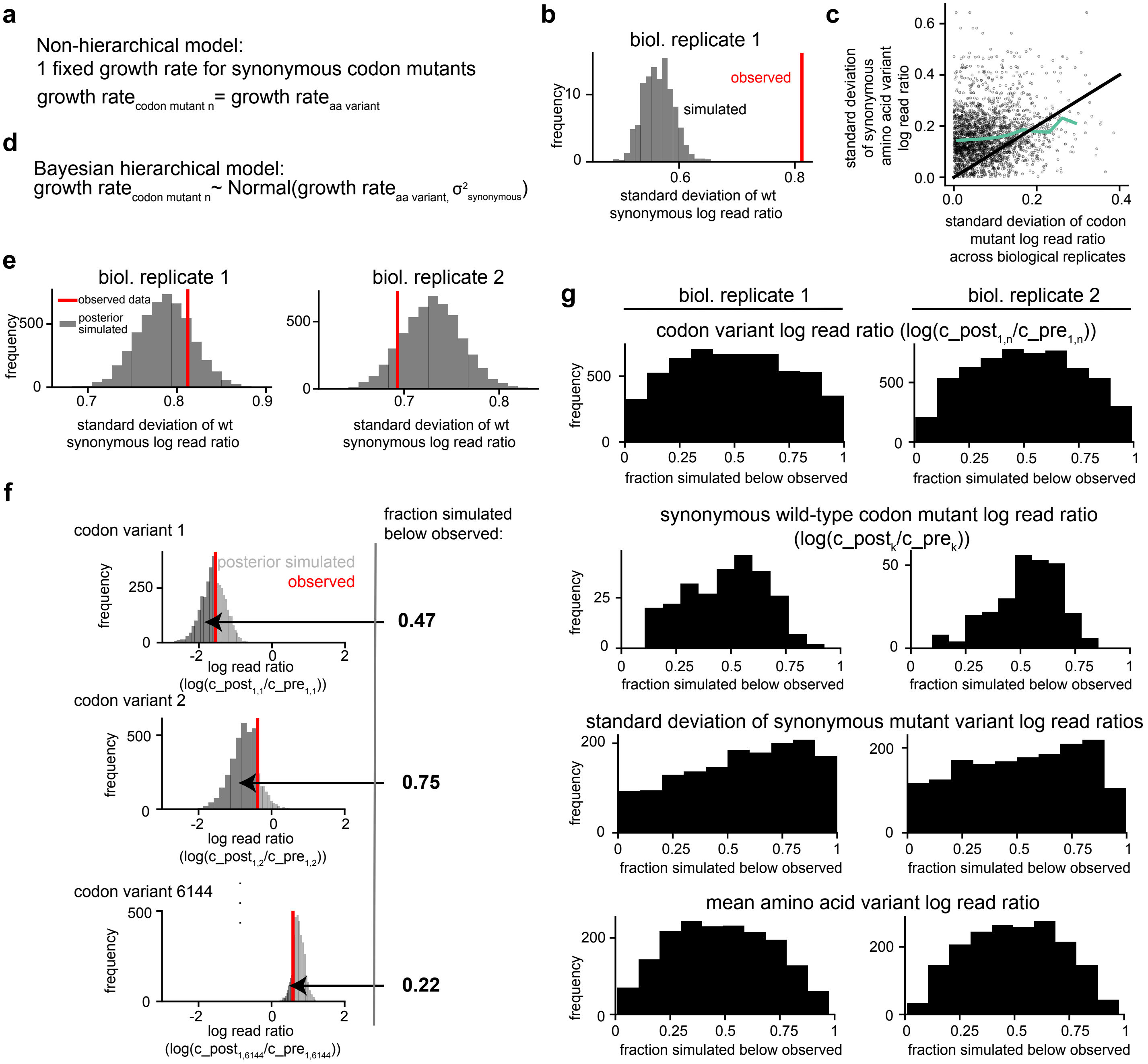


**Extended Data Figure 21:** Posterior predictive checks show that the Bayesian hierarchical model can capture observed data statistics for both replicate experiments, whereas a non-hierarchical model cannot.

**a,b**, A non-hierarchical model, in which all synonymous codon variants have the same growth rate (**a**), cannot explain the observed data. (**b**) The observed standard deviation of log read ratios for synonymous wild-type toxin codon variants (red) (n=278) fall outside of the non-hierarchical model's expectations (grey).

**c**, The synonymous amino acid mutant standard deviations within a replicate (y-axis) are higher than codon mutant standard deviations between replicates (x-axis). Light green indicates binned average.

**d**, Bayesian hierarchical model allows for growth rate variation between synonymous codon mutants by drawing these from a Gaussian distribution.

**e-g**, Observed data statistics fall within the hierarchical Bayesian model's expected values. (**e**) The observed standard deviation of synonymous wild-type toxin codon mutant log read ratios (red) fall within the model simulated values (stdev(log(c_post1$_k$/c_pre1$_k$) or stdev(log(c_post2$_k$/c_pre2$_k$) for biological replicate 1 or 2 respectively), see model code). Compare to panel (b) for the non-hierarchical model. (**f**) For each codon mutant, the hierarchical Bayesian model allows for simulating pre- and post-selection read counts (log(c_post$_{i,n}$/c_pre$_{i,n}$), see ED Fig. 21), including log read ratios, using the posterior parameter distribution. For each codon mutant, we calculate the p-value statistic (ie. the fraction of simulated samples falling below the observed log read ratio). (**g**) Distribution of posterior simulated p-values for various statistics, demonstrating that no observed data statistic is biased to fall outside of the posterior simulated statistics.

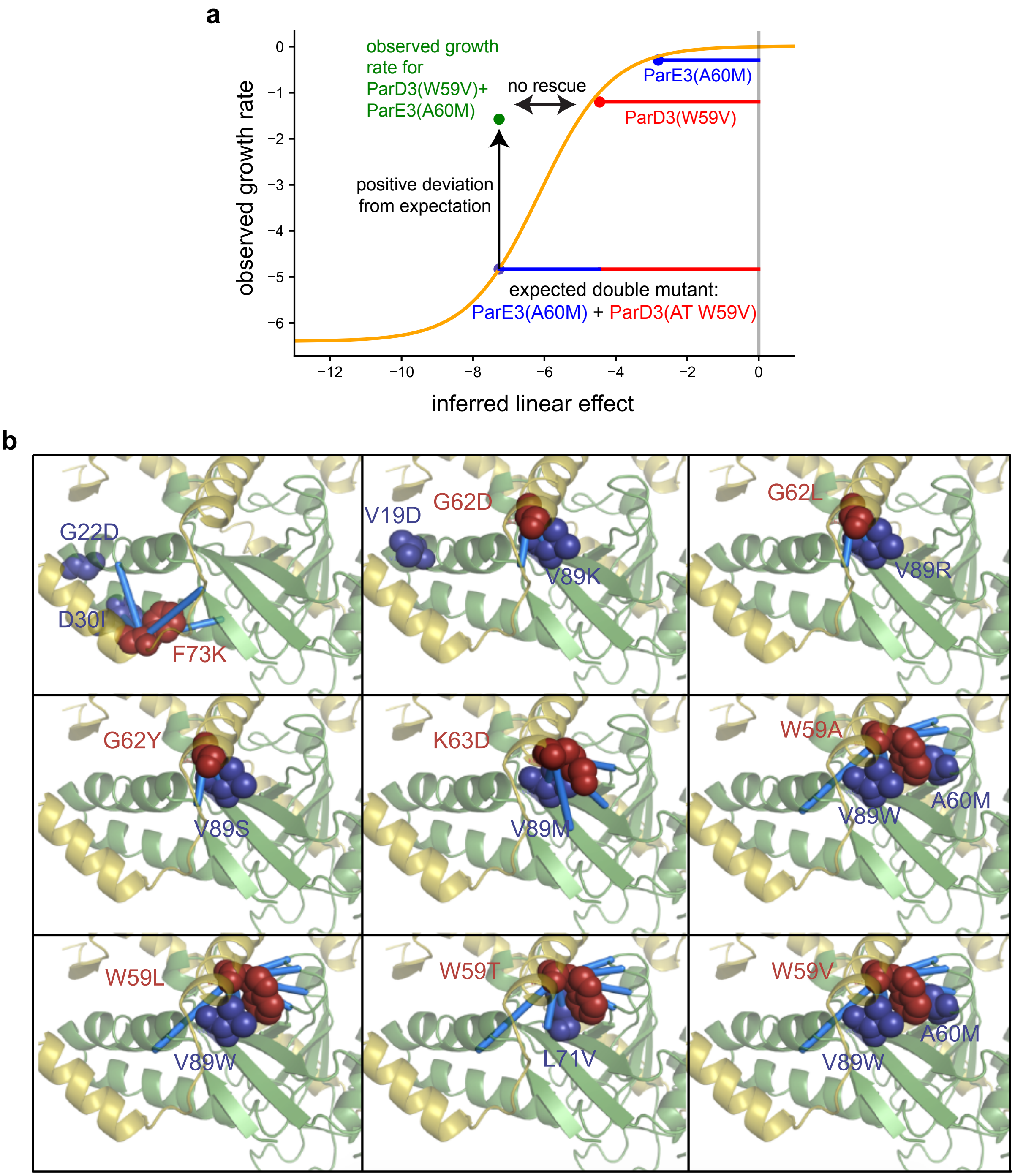


**Extended Data Figure 22:** Toxin-antitoxin double mutant growth rates that deviate positively from the model-predicted expected growth rates do not necessarily rescue the deleterious antitoxin effect, but are mostly physically close.

**a**, Plot indicating how the observed growth rate for the combination of ParE3(A60M) and ParD3(W59V) positively deviates relative to the model-based expected double mutant growth rate, but does not improve the growth rate relative to the antitoxin single mutant alone.

**b**, Almost all of the most positively deviating double mutant residue pairs (T*/AT*) are in direct contact, as indicated on the wild-type toxin-antitoxin crystal structure (PDB ID: 5CEG). Red: deleterious antitoxin, purple: top two postively deviating toxin variant residues, blue dashes: coevolving residues in natural sequences.

### Materials and Methods

#### Bacterial strains, vectors and media

*E. coli* strains were grown at 37 °C in M9L medium (1x M9 salts, 100 μM $CaCl_2$, 0.4% glycerol, 0.1% casamino acids, 2 mM $MgSO_4$, 10% v/v LB). Antibiotics were used as follows: 50 μg/ml carbenicillin, 20 μg/ml chloramphenicol in liquid media, and 100 μg/ml carbenicillin, 30 μg/ml in agar plates. The toxins (ParE3 or ParE2) were carried as before[8] on the pBAD33 vector (chlor$^R$ marker, ML3482 for wild-type ParE3, ML3303 for wild-type ParE2) with expression repressed or induced with 1% glucose and L-arabinose at indicated concentrations, respectively, and the antitoxin ParD3 was carried on the pEXT20 vector (carb$^R$ marker, ML3483) with expression induced by IPTG[1]. Toxin and antitoxin libraries containing all possible single mutants were each cloned under a bicistron RBS design, in which a short leader peptide is engineered upstream of, and co-operonic with, the toxin or antitoxin. This design substantially reduces expression effects that would otherwise arise by variant 5' regions of the toxin or antitoxin forming secondary structure with its ribosome binding site[2,3]. Consistent with this desired effect, we confirmed that synonymous variants throughout the toxin and antitoxin behaved comparably (Extended Data Fig. 18).

For the single mutant and suppressor scan, we used the arabinose-titratable wild-type *E. coli* strain BW27783[4]. For the combinatorial antitoxin library experiments, we used the previously optimized TOP10 *E. coli* background[1].

#### Library construction

The toxin and antitoxin single mutant libraries were each constructed using a previously described 2-step overlap-extension PCR protocol[5]. For the toxin library, we used pBAD33-*parE3* as a template. To introduce mutations at a given amino-acid position/codon, we used a pair of mutagenic primers containing NNNs at the position to be mutated (forward and complementary reverse mutagenesis primers; see Table S2). The reverse mutagenesis primer was used with the primer DDP115 specific to the 5' end of *parE3* and the complementary forward mutagenesis primer was used with the primer DDP116 specific to the 3' end of *parE3* (PCR cycling was: 30 sec. at 98°C; 20 cycles of:10 sec. at 98°C, 20 sec. at 55°C, 1 min. at 72°C; 2 min. at 72°C, hold at 4°C; using Phusion kit (NEB) or KAPA). The products of these two PCRs were then combined, diluted 1:100, and amplified with DDP115 and

DDP116 using the same thermocycling protocol to create full-length *parE3* harboring all possible nucleotides at a single codon position. For codon positions that failed to yield a desired, full-length PCR product on a 1% agar gel, we added 3% DMSO, 1M betaine and/or 6% 1,2-propanediol. This same process was repeated for each possible position in *parE3* and the final PCR products combined in approximately equimolar concentrations using the Qubit kit (ThermoFischer). The same overall process was followed to create the antitoxin mutant library, but used DDP141 and DDP142 as the flanking primers and pEXT20-*parD3* as the template. We then amplified the pBAD33-*parE3* and pEXT20-*parD3* vectors using primers DDP508 and 509 or DDP540 and 541 (following KAPA kit thermocycling recommendations), respectively. These PCR products were digested with HindIII-HF and SacI-HF (NEB), and then subjected to a PCR clean up kit (Qiagen or Zymo). The PCR products that comprised the toxin and antitoxin library inserts were also digested with HindIII-HF and SacI-HF, subjected to a PCR clean up kit (Select-a-size DNA Clean&Concentrate with >150 base-pair size cutoff to manufacturer specifications), and then ligated into the amplified vectors using T4 DNA ligase (NEB) at 16 ºC for 16 hours with a 1:3 molar ratio of insert to vector with 50 ng vector per 20 µl ligation reactions. These ligation reactions were scaled based on downstream needs. Ligations were dialyzed on Millipore VSWP 0.025 µm membrane filters for 90 minutes before transformation.

**Single mutant deep mutational scanning library preparations**

For the antitoxin single mutant library, we transformed in replicate ~40 µl of electrocompetent BW27783 (made electrocompetent as described previously[6]) harboring the wild-type toxin on pBAD33 with 1-2 µg of dialyzed antitoxin single mutant ligation reactions, recovered in 1 ml SOC for 1 hour at 37 ºC. We plated 10 µl of the recovered transformants in a 1:10 serial dilution series on selective agar plates (carbenicillin/chloramphenicol/1% w/v glucose) to check transformation efficiency (all libraries were propagated with $>10^6$ transformants). We minimized the number of generations that the libraries were propagated within cells due to leaky toxin expression, and grew cells at 37 °C until they reached $OD_{600}$ ~ 0.3, at which point glucose was removed by washing 4 times with M9L (spinning cells down at 8000 *g* for 5 min), and ready for the growth rate measurements described below.

For the toxin single mutant library in the presence of wild-type antitoxin, we followed the process outlined above for the antitoxin library with the exception that we transformed electrocompetent BW27783 cells containing wild-type antitoxin on pEXT20 with the toxin single mutant ligation reactions. For the toxin single mutant library in the absence of antitoxin, we transformed electrocompetent BW27783 cells containing an empty pEXT20 plasmid with the toxin single mutant ligation reactions, and followed the same process as above.

**Suppressor library preparation**

For the suppressor scan, we screened all toxin single substitutions against a set of 36 antitoxin single substitutions. 29 of the 36 antitoxin substitutions are found at positions that are within the top 15 toxin-antitoxin covarying pairs of residues with the toxin, 5 of the 36 are found within the top 7 antitoxin-antitoxin covarying pairs of positions (E26, I33, W41, V17, K44), and 2 did not fall in either of these categories (A16K, D60I). For the suppressor scan, we measured the bulk growth of cells pooled in a single flask containing the toxin single mutant library plasmids with up to 15 different antitoxin mutant backgrounds (see github repository for pooling of variants). To read out both the toxin single mutant and antitoxin single mutant background by sequencing the toxin gene only, we cloned the toxin mutant library into separate vectors containing one of 15 different 4-nucleotide barcodes just 3' to the toxin gene and restriction site. Barcoded vector backbones of pBAD33 were generated using primers (reverse primer DDP 508 and forward primers DDP239-254, using KAPA kit and KAPA recommended cycling protocols) that contain and therefore introduce the barcodes. Barcodes were chosen to be at least 3 nucleotides different from any other barcode (Table S2). These barcoded vector backbones were then separately ligated with the toxin single mutant library insert and prepared as described above. We transformed this DNA ligation reactions separately into TOP10 cells, grew them to $OD_{600}$ ~ 0.5 at 37 °C in M9L/carb/chlor/1% glucose and then miniprepped plasmids. We then transformed each barcoded toxin library in replicate into electrocompetent BW27783 cells containing one of these antitoxin single mutant plasmids, and kept track of which barcode corresponds to which antitoxin single mutant. The antitoxin single mutant plasmids were generated by following the above overlap extension PCR protocol with mutagenesis primers described in Table S2 and flanking primers DDP141 and DDP142 using the pEXT20-*parD3* plasmid as template. This allowed us to sequence the full-length toxin gene and adjacent 10 nucleotides to determine both the toxin mutation, as well as the antitoxin mutant background from pooled cell samples.

Each of the transformed libraries was recovered in 1 ml SOC, transformation efficiency checked as described above, and then grown in 50 ml M9L/carb/chlor/1% glucose to $OD_{600}$ ~ 0.3, resuspended in

5 ml M9L/carb/chlor/1% glucose/20% glycerol, aliquoted into 1 ml tubes, and flash frozen in liquid nitrogen. On the day of growth rate measurements, aliquots of the toxin single mutant libraries in each of the different antitoxin backgrounds were thawed, equally mixed based on their $OD_{600}$ measurement at the time of freezing, and recovered in 50 ml M9L/carb/chlor/1% glucose at 30 °C for 3 hours. Subsequently, glucose was removed by washing 4 times with M9L, and cells were ready for growth rate measurement.

**Combinatorial antitoxin library preparation**

For the combinatorial mutant scanning, we followed the previously optimized expression conditions[1] under which wild-type ParD3 shows differential neutralization of cognate and non-cognate toxins. TOP10 *E. coli* cells containing the combinatorial library were made electrocompetent[6]. A plasmid containing wild-type ParE3, wild-type ParE2, or one of 10 ParE3 variants, generated using overlap-extension PCR[5] (as above using wild-type toxin plasmid as template and mutagenesis primers DDP627-642), was then transformed into these cells, transformation efficiency checked, and cells grown up in 50 ml M9L/carb/chlor/1% glucose for snap-freezing. Cells were then thawed, recovered at 30 °C for 3 hours and glucose washed off as above.

**Library growth rate measurements**

On the day of growth rate measurement, cells prepared as detailed above were resuspended at an $OD_{600}$ of ~0.03 in 250 ml M9L/carb/chlor/IPTG, which induces antitoxin expression. After 100 minutes, arabinose was added to induce toxin expression (t=0 minutes). Cell growth was followed by measuring $OD_{600}$ every 30-60 minutes, and when cultures reached an $OD_{600}$ of ~0.3, they were diluted 1:10 with fresh, pre-warmed M9L/carb/chlor/IPTG/arabinose to keep cells in exponential growth throughout the duration of the experiment. Samples were taken at the time of toxin induction, and 10 hours after toxin induction, which corresponds to ~6 generations of cells harboring wild-type ParE3 and ParD3. This results in a dynamic range of $2^6$ = 64-fold change. Each library growth experiment was performed in duplicate, using separate library transformations for each replicate.

For the single mutant and suppressor scans, performed in the arabinose titratable strain BW27783, toxin was induced by adding the lowest concentration of arabinose (1.2 x $10^{-4}$ %) that produced complete cessation of growth using the wild-type toxin (Extended Data Fig. 5b). For this concentration of arabinose, 10 mM IPTG was the minimal concentration necessary for the wild-type antitoxin to provide a nearly complete restoration of growth (Extended Data Fig. 5c). To assess whether our results are sensitive to the chosen induction levels, we performed the entire suppressor scan (73,440 amino acid variants) also at a 'low' antitoxin concentration that almost, but not fully, neutralizes the toxin (8 x $10^{-5}$ % arabinose, 17.5 mM IPTG, Extended Data Fig. 5c). For the toxin single mutant scan done in the absence of wild-type antitoxin, we performed the growth measurement at 7 different arabinose induction levels (0.2, 5.3 x $10^{-3}$, 8 x $10^{-4}$, 1.2 x $10^{-4}$, 8 x $10^{-5}$, 4 x $10^{-5}$, 2 x $10^{-5}$ % arabinose) for which the highest 4 reach full growth inhibition (Extended Data Fig. 4b,c), each in replicate. For the combinatorial mutant scans, we performed the experiments with previously optimized inducer concentrations of 0.2% arabinose (w/v) and 100 μM IPTG induction in the TOP10 strain background.

**Sample sequencing and preparation**

For each replicate, samples taken at 0 and 600 minutes after toxin induction were miniprepped (Zymo Research Plasmid Miniprep Kit). We then performed a high-input (400 ng plasmid DNA), low cycle (12 rounds) PCR reaction using KAPA-HiFi (cycling conditions) to amplify the toxin or antitoxin library of interest. The primers introduced Illumina adapter sequences, sequencing primer homology regions, and Illumina multiplex indices for each sample (forward primer DDP543 and reverse primers DDP178-193 + DDP569-580 for the toxin in single mutant and suppressor screens, forward primer DDP544 and reverse primers DDP545-568 for antitoxin single mutant library, forward primers DDP643-645 with reverse primers DDP646-693 for combinatorial antitoxin library). These primers also introduce variable numbers of random nucleotide or YRYR nucleotides (Y corresponding to random pyrimidines, R corresponding to random purines) as the very first bases to be sequenced on the forward primers in order to allow for Illumina cluster definition, and stagger our homopolymer-like amplicon.

We gel purified amplicons of interest (~500 nucleotides) by running samples with a loading dye for 30 minutes at 180 V on a Novex 8% TBE gel. We sheared the excised gel band by spinning it through a bottom-pierced 0.5 ml tube placed within a 1.5 ml tube, added 500 μl of 10 mM Tris buffer (pH=8), and then froze the sample at -20 °C for 15 minutes followed by incubation at 70 °C for 10 minutes to solvate the DNA from the gel. We spun each sample through a 0.22 μm spin-x cellulose acetate column to separate the gel from the supernatant, and then performed an isopropanol precipitation of the DNA by adding 32 µl 5M NaCl, 2 µl glycoblue, and 550 µl 100% isopropanol. The mixture was then chilled at -80 °C and centrifuged at 4 °C at 14,000 *g*. Finally, the sample was washed with ice cold 70% ethanol, air-dried, and resuspended in 10 µl water.

Each sample was then run on a fragment analyzer, and qPCR was used to quantify the DNA concentration. Finally, samples were pooled and sequenced on a MiSeq, or HiSeq 2500, with 250 or 300 bp paired end reads. Sequencing was performed with variable 20-30% PhiX spike in.

**Analysis of sequencing data and growth rate measurements**

Raw fastq paired-end sequencing reads were merged using FLASH 1.2.11[7]. Merged reads were quality filtered based on their phred-score using vsearch 2.13.0[8], with the following arguments: vsearch --fastq_filter {0} --fastq_truncqual 20 --fastq_maxns 3 --fastq_maxee 0.5 --fastq_ascii 33 --fastaout {1}.fasta

Toxin mutant reads were subsequently split into separate files based on their four-nucleotide barcode indicating the antitoxin background, if applicable. Subsequently, reads were filtered for those that (i) spanned the full-length of the toxin or antitoxin gene, (ii) had no mutations or indels in the immediately flanking 10 bp upstream (which includes the restriction site, RBS, and the stop codon for the upstream bicistronic peptide), as well as the downstream 6 bp (which includes the restriction cloning site). The majority of reads (~77%) contained a single codon mutation. Custom analysis scripts for raw read processing, Bayesian inference of growth rates (see below) and nonlinear modeling (see below) are available at: https://github.com/ddingding/coevolution_paper.

**Hierarchical Bayesian inference of mutant growth rates**

We used a Bayesian model (Extended Data Fig. 19) that allowed us, given a plausible data generating process (likelihood function) that captures how growth rates could give rise to our observed codon-level read count data before and after selection, and vague priors, to get posterior probabilities for our growth rates of interest, namely how likely different values of a growth rate for a particular amino acid substitution are given the data observed. This model takes into account sampling noise of reads, synonymous mutant observations per amino acid, and biological replicate experiments to infer amino-acid variant growth rates and their uncertainties. Our model allowed for calibrated uncertainty inference, as well as unbiased inference of amino acid substitution effects compared to the widely-used log-read ratio statistic (Extended Data Fig. 20).

For our likelihood model, we extended previous Bayesian approaches for mutant effect inference from read data[9,10] and built a hierarchical Bayesian model. As done previously, read counts for each codon at a particular time-point were emitted from a Poisson distribution with an inferred Poisson rate parameter λ. The post-selection Poisson rate parameter is the pre-selection Poisson rate multiplied by the exponentiated growth rate for each codon mutant (following exponential growth rates of the form $N_t = N_0 * e^{(\text{growth rate*t})}$).

We found that previously used models, in which all synonymous codon variants share the exact same growth rate, were insufficient to explain the observed variance in read ratios for synonymous variants (Extended Data Fig. 21a-c). This finding motivated us to expand from a non-hierarchical generative model to a multilevel model allowing for partial pooling of synonymous variant growth rates to inform inference of their shared, amino-acid-level variant growth rates. For this model, the growth rate for a particular codon mutant was drawn from a normal distribution whose mean is the growth rate of the amino-acid variant. In this way, each synonymous mutant's growth rate informs the amino-acid variant growth rate, while still capturing the observed variation in synonymous mutant growth rates.

Because the posterior of our model is not analytically tractable, we used Stan[11,12] to perform inference (using 2 MCMC chains, 10,000 steps each, discarding the first half of MCMC chains as 'warmup'). This gave use 10,000 discrete samples for each amino-acid variant growth rate of interest, approximating the true continuous posterior density. We used the mean of these 10,000 samples as our best guess for the true growth rate of that particular amino-acid variant, with the distribution of these 10,000 samples reflecting the posterior uncertainty in the inferred growth rate.

**Growth rate inference validation**

We validated our Bayesian model in three ways, as summarized below.

*Inference of synthetic growth rates*: We generated three different synthetic datasets (pre- and post-selection read counts for each codon variant) by assuming three different distributions of true amino-acid mutant growth rates (from the range of antitoxin single mutant growth rate distributions). We compared the Bayesian inference to the classically used log read ratio (+1 pseudo-count), and found that our model removed the bias introduced in the raw log read ratio when few reads are observed post-selection (Extended Data Fig. 20). The mean posterior growth rate for each amino acid mutant still correlated well with the true synthetic growth rates at low growth rates. As desired, our model assigns these low growth rate values with few observed post-selection reads a larger 95% highest posterior density interval.

*Calibrated uncertainty*: Our Bayesian model allowed us to calculate the associated uncertainty, *i.e.* the posterior distribution for each amino-acid variant. We compared differing uncertainty intervals for each variant (95%, 90%, 80%, 50% posterior highest density intervals) with the true growth rates used to simulate the observed count data, and found that the percentage of true growth rates falling within the posterior highest density intervals corresponded to the percentage of the highest density interval (Extended Data Fig. 20d).

*Posterior predictive checks*: Posterior predictive checks allowed us to assess whether a given model was complex enough to capture the observed data. After model parameter inference, replicate data were generated from the model using the inferred distribution of parameters, and compared to the observed data (Extended Data Fig. 21d-g). We chose multiple different test quantities (log read ratios for each codon and averaged across amino-acid variants, the standard deviation of synonymous mutant log read ratios for each amino-acid variant as well as synonymous wild-type toxin mutants) to compare 10,000 simulated replicate datasets generated from the model to the true observed data, and for each quantity calculated their posterior predictive p-value (*i.e.* the fraction of simulated data above the observed data along the test statistic). Based on these test quantities, the model developed plausibly generates the observed data, demonstrating sufficient complexity.

**Calling significantly beneficial toxin mutants**

Toxin mutants were called as significantly beneficial to a given antitoxin mutation if they grew at least 0.5 $\log_2$-fold better than that antitoxin mutant combined with the wild-type toxin measured in the same flask, with all 10,000 posterior samples exceeding the growth rate of the antitoxin mutant with the wild-type toxin, i.e. the 99.99% highest posterior density interval does not overlap zero difference in growth rate. For the ‘high' antitoxin expression condition (see main text), we sought beneficial toxins in 9 deleterious antitoxin backgrounds (F73K, K63D, W59A/L/V/T, G62D/Y/L). For the ‘low' antitoxin expression condition, we sought beneficial toxins in 12 deleterious antitoxin backgrounds (the former plus E26R, E79H, E79K).

We verified that the toxins harboring beneficial mutations were still as, or nearly as, toxic as the wild-type toxin by measuring the growth rate of the toxin single mutant library at 7 different toxin induction levels as indicated above (Extended Data Fig. 4). For the 2,040 possible amino acid substitutions in the toxin, we called 310 as fully toxic as they did not show any significant growth rate differences (the 95% highest posterior density interval overlapped with the growth rate of the wild-type across all 7 expression conditions. Individual, low-throughput validation of four such toxin mutants revealed no distinguishable growth rate differences to the wild-type toxin (Extended Data Fig. 14) at both the minimum expression level under which the wild-type toxin produces maximal growth inhibition (1.2 x $10^{-4}$% arabinose) in the absence of an antitoxin, as well as when reduced to a level that produced half-maximal inhibition with the wild-type toxin (6 x $10^{-5}$% arabinose). We also called as toxic a less stringently defined set of 781 mutants that were not significantly different from the wild-type toxin across the four highest expression conditions, which are conditions that lead to full growth inhibition by the wild-type toxin (Extended Data Fig. 4).

**Combinatorial mutant scan analysis**

Following previous analyses[1,13], we calculated for each antitoxin amino-acid variant (combining synonymous mutants) a mean log-read ratio for 600 and 0 minutes post-induction of the toxin, with a pseudo-count added to both time-points. We then scaled the growth rates to between 0 and 1 based on the mean growth rate of truncated antitoxin mutations and the wild-type antitoxin, respectively. In the case of the antitoxin library in the background of the non-cognate toxin ParE2, we used the rewired antitoxin ParD3 ILK for the maximum growth scaling.

**Nonlinear modeling and calling of specific vs. non-specific**

We pooled all of the observed growth rate data for toxin mutants combined with the wild-type antitoxin, one of the antitoxin single mutants, or no antitoxin. The predictor for each growth rate is the one-hot encoded amino acid sequence, indicating the site-wise presence of a particular toxin substitution or antitoxin substitution at each position. We fit weights associated with each unobserved, single substitution effect, passed through a global sigmoid function by using the Adam optimizer[14] to minimize the mean squared error of predicted growth rates relative to the observed growth rates. This was done using Tensorflow 2[15], run for 4,000 steps until no decreases in fitting accuracy were observed. Weights were initialized with the ordinary least square weights and Adam learning rates were grid-scanned among [1, 0.1, 0.01, 0.001]. The linear fitting was done similarly using the Statsmodels python package[16], with an identical model missing the sigmoid transformation.

Generally speaking, relative growth rate comparisons should be most robust when comparing mutants grown in the same flask because a particular mutant with one fixed absolute growth rate might decrease in relative fraction in a flask with faster growing mutants, but increase in a flask with slower growing mutants. In our case, the distribution of growth rates in each flask were similar enough, such that a nonlinear model trained on relative growth rates pooled across flasks could fit and explain 89% of the observed growth rate variance, giving us confidence that this issue is a negligible factor in our analysis.

To call a particular double mutant combination as having an observed growth rate that deviates substantially and significantly from that expected by the model, we required the double mutant growth rate to be at least 2-fold greater than or less than the independent expectation, with a 99.99% highest posterior density interval that did not overlap the independent expectation (*i.e.*, all 10,000 samples were greater or less than the expectation). The most positively deviating toxin mutation in each deleterious antitoxin background were mostly in direct contact and were biochemically rationalizable (Extended Data Fig. 22b). However, most of these positively deviating double substitutions did not improve the growth rate over the antitoxin single substitution growth rate, but deviate because their independent expectations are highly detrimental (Extended Data Fig. 22a).

In contrast to previous studies[17–26], we demonstrated that the inferred, unobserved mutation effects in our nonlinear model were robust with respect to the details of expression conditions. In particular, the inferred mutant effects for all toxin mutants were highly correlated for the 'high' and 'low' induction levels of antitoxin (r=0.98, Extended Data Fig. 9h).

**Low-throughput toxicity measurement and orthogonal growth rate validation**

To assess whether non-specific suppressor mutations in the toxin maintained toxicity, we measured the growth rate of cells containing these toxin mutations in the absence of the antitoxin in the arabinose titratable strain at both full (1.2 x $10^{-4}$% arabinose) and half-maximal induction levels (6 x $10^{-5}$% arabinose). We diluted saturated overnight cultures 1:400 into the wells of a 96-well plate containing

M9L supplemented with 10 µM IPTG and arabinose (concentration as indicated, Extended Data Fig. 14), as well as carbenicillin and chloramphenicol. We ran a maximum of 8 samples per 96-well plate, such that each sample is measured at least 10 times in each plate. Row A was kept blank. Each sample was staggered diagonally across the plate reader to minimize plate reader position biases. For example, sample 1 was loaded into wells B1, C2, D3, E4, F5, G6 H7, B9, C10, D11, E12. Each plate also contained samples corresponding to a wild-type toxin combined with a wild-type antitoxin and the wild-type toxin combined with an empty vector for within plate growth rate comparisons. We used the plate reader Biotek Synergy H1 with orbital shaking 365 rpm at 37 ºC, with 180 µl of media and 70 µl mineral oil on top to prevent evaporation.

For orthogonal growth rate validations, we diluted overnight cultures 1:50 into 10 ml of fresh M9L/carb/chlor/1% glucose/10 µM IPTG to pre-induce antitoxin, grew cells 2-3 hours at 37 ºC to $OD_{600}$~0.5, then washed 4 times with M9L. We then diluted these cells 1:200 into 96-well plates containing M9L/carb/chlor/10 µM IPTG/1.2 x $10^{-4}$ % arabinose to measure growth rates. We calculated the growth rate as the normalized fold change in $OD_{600}$ across time. Error bars in Extended Data Fig. 1a are calculated using the error propagation formula for independent variables and a first order Taylor expansion.

**Coevolution analysis**

We performed covariation analysis similar to before[27]. Briefly, we generated JackHMMR[28] alignments using our wild-type toxin (ParE3) and antitoxin (ParD3) as query sequences at a range of bitscore cutoffs (ranging from 0.1 to 0.9) from the uniref100 database. For each bitscore, we concatenated toxin and antitoxin sequences based on their genome distance (< 1000 nucleotides). We selected the bitscore cutoff with the highest number of true positive (minimum atom distance < 6Å) between-protein covarying pairs, resulting in the bitscore choice of 0.3. Alignment quality filtering was done similar to previous studies, calculating covariation scores only for residue positions with at least 80% coverage across sequences, discarding sequences if they did not span 80% of the full-length concatenated query sequence (196 amino acids), and down-weighting sequences if their sequence identity exceeded 80%. The final alignment contained 1650 concatenated sequences, with an effective number of sequences of 1088 after down-weighting.

We chose to highlight the top 10 covarying residues from this analysis, which were all close (< 6 Å minimum atom distance) in distance. Using previous calibration sets[29], the 90% precision cutoff (pairs of residues < 6Å distance/all selected pairs) in covariation score resulted in 29 covarying pairs of residues between protein, of which 28 are indeed within 6 Å minimum atom distance.

For covariation score comparison against other complexes (Fig. 3b), we looked at the top N (with N being 1, 2, 3, 4, 5, or 10) covariation scores between proteins, compared against the corresponding top N covariation scores for other complexes from ref[29].

**Network visualizations**

Force-directed graphs were constructed using the python package network[30]. Network clusters were defined using Louvain clustering. Sequence motifs were visualized for each cluster using Logomaker[31].

**EVmutations and VAE prediction calculations**

EVmutations and VAE predictions were performed as previously described[27,32].

**Structural analysis**

We used the wild-type toxin ParE3-ParD3 (PDB: 5CEG) for structural analysis and distance calculations. Minimum atom distances refer to the minimal atom centroid distance between any two atoms in two different amino acids. Minimum atom distance of a toxin residue to any antitoxin atom were calculated for the minimum of particular toxin residue from toxin chain B or chain D in PDB:5CEG.

**Table S1. Spatial distances of rescuing toxin substitutions to the antitoxin**

| Classification | Toxin variant | Min. atom distance to deleterious antitoxin [Å] | Close to deleterious antitoxin residue (<6Å) | Min. atom distance to any at residue [Å] | Close to any antitoxin residue (<6Å)? | Notes |
|---|---|---|---|---|---|---|
| nonspecific | A12T | 12.23 | 0 | 6.46 | 0 | may contact ParD3 positions E79 & K83 |
| nonspecific | V5Y | 10.22 | 0 | 3.48 | 1 | contacts ParD3 at positions 69-71 on the interface |
| nonspecific | V5M | 10.22 | 0 | 3.48 | 1 | contacts ParD3 at positions 69-71 on the interface |
| nonspecific | V5L | 9.84 | 0 | 3.48 | 1 | contacts ParD3 at positions 69-71 on the interface |
| nonspecific | A66F | 16.35 | 0 | 6.45 | 0 | may contact ParD3 residues E37 and E27, or contribute to toxin dimerization |
| nonspecific | A66Y | 19.94 | 0 | 6.45 | 0 | may contact ParD3 residues E37 and E27, or contribute to toxin dimerization |
| nonspecific | A66P | 19.94 | 0 | 6.45 | 0 | might contribute to toxin dimerization |
| nonspecific | D55E | 14.68 | 0 | 2.66 | 1 | might strengthen existing salt bridge |
| nonspecific | M20I | 17.28 | 0 | 3.41 | 1 | might improve interface complementarity |
| nonspecific | M20L | 17.41 | 0 | 3.41 | 1 | might improve interface complementarity |
| nonspecific | R27E | 26.34 | 0 | 3.50 | 1 | may form new salt bridge with ParD3(R82) |
| nonspecific | R27S | 26.34 | 0 | 3.50 | 1 | may form new H-bond with ParD3(R82) |
| nonspecific | E84Q | 16.90 | 0 | 10.44 | 0 | likely does not make new contact |
| nonspecific | V75K | 12.26 | 0 | 6.25 | 0 | likely does not make new contact |
| nonspecific | R100Q | 18.42 | 0 | 2.82 | 1 | may increase contact or promote dimerization |
| nonspecific | E87Q | 4.70 | 1 | 2.96 | 1 | contacts ParD3 at K63 |
| nonspecific | S23L | 24.60 | 0 | 3.14 | 1 | contacts ParD3 at L84 and K5 |
| nonspecific | A38Y | 20.60 | 0 | 6.28 | 0 | unlikely to make new contact, ~6 Å to ParD3(F73), but faces in opposite direction |
| nonspecific | P54R | 15.57 | 0 | 5.96 | 0 | likely does not make new contact |
| nonspecific | S7T | 6.71 | 0 | 3.55 | 1 | contacts ParD3 at G62 |
| nonspecific | V83L | 17.46 | 0 | 12.52 | 0 | not close to any ParD3 residue |
| specific | E87H | 4.70 | 1 | 2.96 | 1 | contacts ParD3 P68 and likely contacts to ParD3 G62 |
| specific | E87K | 4.70 | 1 | 2.96 | 1 | contacts ParD3 P68 and likely contacts to ParD3 G62 |
| specific | T9L | 6.87 | 0 | 2.95 | 1 | contacts ParD3, but on other side of helix of ParD3 W59 |
| specific | T9C | 6.87 | 0 | 2.95 | 1 | contacts ParD3, but on other side of helix of ParD3 W59 |
| specific | E87N | 4.70 | 1 | 2.96 | 1 | contacts ParD3 P68 and likely contacts to ParD3 G62 |
| specific | R57N | 9.61 | 0 | 2.77 | 1 | contacts ParD3 W59 and D60 |
| specific | A60T | 3.55 | 1 | 3.55 | 1 | contributes to pocket for ParD3 W59 |
| specific | E73K | 7.97 | 0 | 2.50 | 1 | contacts ParD3 W59 and K63 |
| specific | E73Q | 7.97 | 0 | 2.50 | 1 | contacts ParD3 W59 and K63 |
| specific | P8S | 4.90 | 1 | 3.26 | 1 | contacts to ParD3 loop 66-71 |
| specific | S59A | 3.60 | 1 | 3.60 | 1 | contacts ParD3 W59 |

**Table S2 (separate .xlsx file)**

Primers used in this study.

**Table S3 (separate .xlsx file)**

Strains created in this study.

**File S1 (separate .pse file)**

Location of beneficial toxin substitutions on the crystal structure.

### References (Supplemental)

11. Stan Development Team. Stan Modeling Language Users Guide and Reference Manual, 2.26. (2021).

12. Riddell, A., Hartikainen, A. & Carter, M. *PyStan (3.0.0)*. (2021).

13. Lite, T. L. V. *et al.* Uncovering the basis of protein-protein interaction specificity with a combinatorially complete library. *eLife* **9**, 1–57 (2020).

14. Kingma, D. & Ba, J. Adam: A Method for Stochastic Optimization. *Int. Conf. Learn. Represent.* (2014).

15. Abadi, M. *et al.* Tensorflow: A system for large-scale machine learning. in *12th USENIX Symposium on Operating Systems Design and Implementation* 265–283 (2016).

16. Seabold, S. & Perktold, J. statsmodels: Econometric and statistical modeling with python. in *9th Python in Science Conference* (2010).

28. *HMMER, http://hmmer.org/.*

29. Green, A. G. *et al.* Large-scale discovery of protein interactions at residue resolution using co-evolution calculated from genomic sequences. *Nat. Commun.* **12**, 1396 (2021).

30. Hagberg, A. A., Schult, D. A. & Swart, P. J. Exploring Network Structure, Dynamics, and Function using NetworkX. in *Proceedings of the 7th Python in Science Conference* (eds. Varoquaux, G., Vaught, T. & Millman, J.) 11–15 (2008).

31. Tareen, A. & Kinney, J. B. Logomaker: Beautiful sequence logos in Python. *Bioinformatics* **36**, 2272–2274 (2020).

32. Riesselman, A. J., Ingraham, J. B. & Marks, D. S. Deep generative models of genetic variation capture the effects of mutations. *Nat. Methods* **15**, 816–822 (2018).
